# STAT3-mediated allelic imbalance of novel genetic variant rs1047643 and B cell specific super-enhancer in association with systemic lupus erythematosus

**DOI:** 10.1101/2021.09.20.461037

**Authors:** Yanfeng Zhang, Kenneth Day, Devin M. Absher

## Abstract

Mapping of allelic imbalance (AI) at heterozygous loci has the potential to establish links between genetic risk for disease and biological function. Leveraging multi-omics data for AI analysis and functional annotation, we discovered a novel functional risk variant rs1047643 at 8p23 in association with SLE. This variant displays dynamic AI of chromatin accessibility and allelic expression on *FDFT1* gene in B cells with SLE. We further found a B-cell restricted super-enhancer (SE) that physically contacts with this SNP-residing locus, an interaction that also appears specifically in B cells. Quantitative analysis of open chromatin and DNA methylation profiles further demonstrated that the SE exhibits aberrant activity in B cell development with SLE. Functional studies identified that STAT3, a master factor associated with autoimmune diseases, directly regulates both the AI of risk variant and the activity of SE in cultured B cells. Our study reveals that STAT3-mediated SE activity and cis-regulatory effects of SNP rs1047643 at 8p23 locus are associated with B cell deregulation in SLE.

## Introduction

Super-enhancers (SEs) are recently discovered large domains of clustered enhancers (1, 2). The extraordinary feature of SEs is the extremely high and broad enrichment of enhancer-related transcription factors (TFs), H3K4me1 and H3K27ac modifications, resulting in high capability to enhance gene expression programs (2). A large quantity of SEs show cell/tissue specificity (3), thereby they have become principal determinants of cell identity (4). Nonetheless, disease-associated SEs, in particular those exhibiting aberrant activity in autoimmune diseases, are less characterized.

Signal transducer and activator of transcription 3 (STAT3), as one of seven STAT family members, is activated by phosphorylation at tyrosine 705 (Y705) and/or at serine 727 (S727) (5). After import to the nucleus, the phospho-STAT3 (pSTAT3) modulates gene transcription by binding its target sequence (6). STAT3 has gained broad attention because it plays a key role in a variety of pathophysiological immune responses related to lymphocyte development and differentiation, and in other cellular processes of normal and tumor cells (7).

Systemic lupus erythematosus (SLE) is an autoimmune disease that is known to be associated with an array of abnormal immune cell signaling. B-cell hyperactivity in auto-antigen recognition and interaction with T-cells, which ultimately results in multi-organ damage, is central to the pathogenesis of SLE (8). Genetic factors conferring a predisposition to the development of SLE have been widely characterized. Over 100 loci have been implicated in SLE by genome-wide association studies (GWAS) (9, 10). Among them, several genes and/or loci are potent as putative drivers of the disease. For example, genetic risk variants at the promoter of *BLK* at 8p23 locus alter *BLK* transcription activity and thus contribute to autoreactive B-cell responses (11). Nonetheless, the GWAS-identified genetic variants together explained approximately 30% of the heritability of SLE (12, 13), suggesting a requirement of further efforts to explain the missing heritability of SLE. Meanwhile, there is growing evidence that genetic risk factors behave in a context-dependent or cell-specific manner (11, 14). Thus, for SLE and other autoimmune diseases, there is a need to identify the regulatory programs in which these genetic factors impact the immune cell developmental processes.

One approach for tying genetic risk to function in the post-GWAS era (14), is a measurement of allelic imbalance (AI) on two alleles at a given heterozygous locus, typically at single nucleotide polymorphism (SNP). The genes and/or loci with SNPs exhibiting AI could provide a strong foundation for implicating the genetic or epigenetic mechanisms linked to complex traits or diseases (15, 16). As a readout of AI, analyses of allele-specific chromatin accessibility and allele-specific RNA expression have accumulated a wealth of interesting findings, including functional cis-regulation (17, 18), genomic imprinting (19), X-chromosome inactivation or escape (20). Therefore, tracking AI difference in a comparison between diseases and controls may enable to uncover novel functional variants associated with complex diseases. In this study, we describe one such strategy through integrative multi-omics analysis to discover known or novel functional variants associated with SLE, and report on the identification of a novel risk variant rs1047643 and B cell specific SE in B cells with SLE. We further demonstrate that the resultant allelic imbalanced variant and SE activity are directly controlled by STAT3, a master TF that plays a critical role in B cell development and highly associates with autoimmune diseases.

## Material and methods

### Reagents and Antibodies

ML115 (Cayman Chemical); S3I-201 (SML0330, Sigma); Phospho-STAT3 (Ser727) antibody (Cat No. PA5-17876, Invitrogen), Anti-Histone H3 (acetyl K27) antibody (ab4729, Abcam), H3K4me1 Recombinant Polyclonal Antibody (Cat No. 710795, Invitrogen), normal rabbit and mouse IgG (Santa Cruz Biotechnology)

### Data collection

We collected a variety of functional genomics data, including ATAC-seq, RNA-seq, RRBS, Hi-C data (see details in Table S1), from the Gene Expression Omnibus (GEO) and ArrayExpress database. Meanwhile, we downloaded genotype and Epidemiological data from a SLE case-control study (accession: phs001025.v1) in Hispanic population (1,393 cases and 8,86 controls) from the dbGaP database with approval (accessed 29 Sep 2020).

### Analysis of RNA-seq and ATAC-seq data

RNA-seq data were analyzed as described previously with few modifications (21). In brief, raw sequencing data were mapped to the human reference genome (hg19) using Hisat2 program (22) with the default setting. Aligned data were processed and converted into BAM files using SAMtools program (23). The fragments per kilobase of exon per million fragments mapped (FPKM) values were estimated from the Cufflinks program to quantify gene expression levels.

We used a similar method described previously with several modifications (24) to process the ATAC-seq data. In brief, raw sequencing data were mapped to the human reference genome (hg19) using Bowtie2 program (25) with the default setting. Tag per million (TPM) metric, a method commonly used for read counting normalization, was used to quantitatively present the enrichment of open chromatin states across regions of interest.

### Identification of allelic chromatin accessibility difference sites

We used a similar approach described previously to call variants and allelic analysis (20). Briefly, the deduplicated reads in BAM format were realigned and recalibrated, and genetic variants were called in a multiple-sample joint manner implemented in the GATK toolkit (version 3.3). We next filtered out variants as follows: (1) mapping quality score < 20, (2) ≥ 3 SNPs detected within 10 bp distance, (3) variant confidence/quality by depth < 2, (4) strand bias score > 50, (5) genotype score < 15 and (6) read depth < 8. Then, we extracted SNPs annotated from dbSNP (Build 150) that were called as heterozygotes for each sample. For a reasonable comparison, those heterozygous SNPs identified at least triple in both case and control samples were retained. Using allelic ratio (20) as a response variable in linear regression model (see below), we conducted AI analysis on chromatin accessibility for each heterozygous SNP in a comparison between SLE and controls.

Allelic ratio ∼ α + β ∗ disease + ε

The p-values and beta coefficients were calculated to estimate the significance of the association, and the differences between cases and controls, respectively.

### Association analysis

For genotype data from a SLE case-control study in Hispanic population, all typed SNPs in chromosome 8 were extracted for imputation using TOPMed Imputation Server (26). To test SNP rs1047643 in association with SLE, we used a method described previously for univariate and haplotype analyses (27). In brief, the per-allele odds ratio (OR) and 95% confidence interval (CI) for the rs1047643 was estimated for SLE risk using a log-additive logistic model with covariates of populations, sex and five principal components (PCs). We used the haplo.stats package in R for haplotype analyses with populations, sex and five PCs as covariates.

### Super-enhancer annotation

We downloaded whole-genome chromatin state segmentation data (core 15-state model) for 127 cell types from the Roadmap project. As Parker et al. (28) defined, we consider contiguous genomic region marked by states 6-7 (enhancer states, annotated by chromHMM) with ≥ 3 kb as SE in a cell type. Then, we extracted and annotated super-enhancers on 8p23 locus.

### Analysis of eQTL data

We collected eQTL data sets from three large-scale studies, the Genotype-Tissue Expression (GTEx, v8) (29), the Haploreg v4.1 dataset (30) and the study by Westra et al. (31). By searching for the SNP rsID or the coordinate, we extracted the linked genes with query SNPs and plotted the results based on the significance and studies.

### Hi-C data analysis

For in situ Hi-C dataset (Accession ID: GSE63525), we downloaded the Hi-C binary file from Rao et al. study (32) and extracted the observed long-range interactions normalized with Knight-Ruiz matrix balancing (KR) method at 10 kb resolution across the 8p23.1 region (the coordinate: chr8:11260000-11740000, hg19). For other genome-wide Hi-C (Accession ID: GSE113405) and capture Hi-C (CHi-C) datasets (Accession ID: GSE81503 and E-MTAB-6621), we used the Hi-C Pipeline (HiCUP) (33) to truncate and align reads to the human reference genome. The deduplicated data were then processed using the Homer pipeline (34) to call the significant chromatin interaction at 10 kb resolution. The resulting interactions were visualized using UCSC Genome Browser or Sushi package in R environment.

### DNA methylation analysis

We downloaded the processed RRBS dataset of DNA methylation profiles on each CpG site from Scharer et al. report (35) , then extracted and compared CpG methylation levels on a region of interest between SLE and healthy controls.

### Cell culture

GM11997 B lymphoblastic (purchased from Coriell Institute) cells were cultured in RPMI-1640 medium, supplemented with 10% FBS (Thermo Fisher Scientific), 2 mM L-glutamine and 1% penicillin-streptomycin at 37 °C with 5% CO_2_. For perturbation of STAT3, B cells were plated in 12-well plates or 10 cm dishes one day prior to the experiment. Cells were then treated with S3I-201 or ML115. Cells were harvested, washed with PBS and analyzed for proper assays.

### Reverse transcription qPCR

Total RNA was isolated from cells using TRIzol Reagent (Invitrogen) according to the manufacturer’s protocol. 1 µg of total RNA was reverse transcribed using SuperScript III reverse transcriptase and random hexamer. One-tenth of the RT reaction was used as a template for real-time PCR using Luna Universal qPCR Master Mix (New England Biolabs) on a QuantStudi 6 system. Relative expression was calculated with 2^−ΔΔCt^ using the average value of housekeeping gene *GAPDH*.

### Chromatin immunoprecipitation

ChIP was performed as described previously. (2) Approximately 10 × 10^6^ suspension cells were harvested and in 10 ml PBS with 1% formaldehyde for 10 min at room temperature, followed by adding 0.125 M glycine for 5 min. Cells were washed and pelleted by centrifugation and lysed with buffer (50 mM Tris-HCl, pH 7.5, 1% IGEPAL CA-630, 1 mM EDTA, 0.1% SDS, plus 1 mM PMSF) in the presence of protease inhibitors and incubated on ice for 30 min. Cell lysate was sonicated to shear DNA to a length of 200–600 bp. The lysates were centrifuged, and supernatant transferred to new tubes. For immunoprecipitation, approximately 2 × 10^6^ cells and 2-3 µg of antibodies or isotype matched IgG as control were used per ChIP and incubated with supernatant at 4°C on a rotating wheel overnight. Chromatin-antibody complexes were sequentially washed with low-salt buffer, high-salt buffer, LiCl buffer, and TE buffer. Cross-links were reversed by addition of 100 µl of 1% SDS plus 100 mM NaHCO_3_ and by heating at 65°C overnight. Following phenol/chloroform/isoamyl alcohol extraction, immunoprecipitated DNA was precipitated with isopropyl alcohol and resuspended in nuclease-free water. For the identification of the specific regions of interest, ∼10 ng of purified DNA was quantified to determine the percentage of each analyzed region against input DNA. The PCR primers are shown in Table S3.

### Statistical analysis

Data were presented as mean ± SD of three replicates unless stated otherwise. Correlation analysis was performed using Pearson’s correlation coefficient. The differences were considered statistically significant at two-sided P-values less than 0.05.

## Results

### Multi-omics data summary

Functional genomics sequencing data sets comprising 279 samples from eleven studies were collected (Table S1). Of eleven studies, seven are SLE case-control studies with data across three immune cell types including B cells, T cells and Neutrophils (Table S2). Also included in the present study were SNP microarray data from a SLE GWAS study (n = 2,279).

### Identification of SLE-associated variant showing AI at both chromatin and RNA levels

We next designed a two-stage study (Figure 1) to identify putative SLE-associated functional variants. In stage I, also termed as the discovery stage, two chromatin accessibility (ATAC-seq) data sets (Accession ID: GSE118253 and GSE71338, Table S1) comprised 49 samples were analyzed. We focused on those variants displaying difference in AI of chromatin accessibility at heterozygous SNP sites in a comparison between SLE and controls (see Methods in detail). From the reciprocal validation between two data sets, SNP rs1047643 was identified to show the significant AI in B cells from patients with SLE, relative to controls (Figure 2A). Interestingly, in B cells at different stages, the allelic preference of chromatin accessibility for this SLE-associated SNP is alterable. For example, the T allele exhibits more preferential chromatin accessibility in activated B cells from patients, relative to the C allele. However, the direction is reversed in SLE naive B cells.

**Figure 1.**
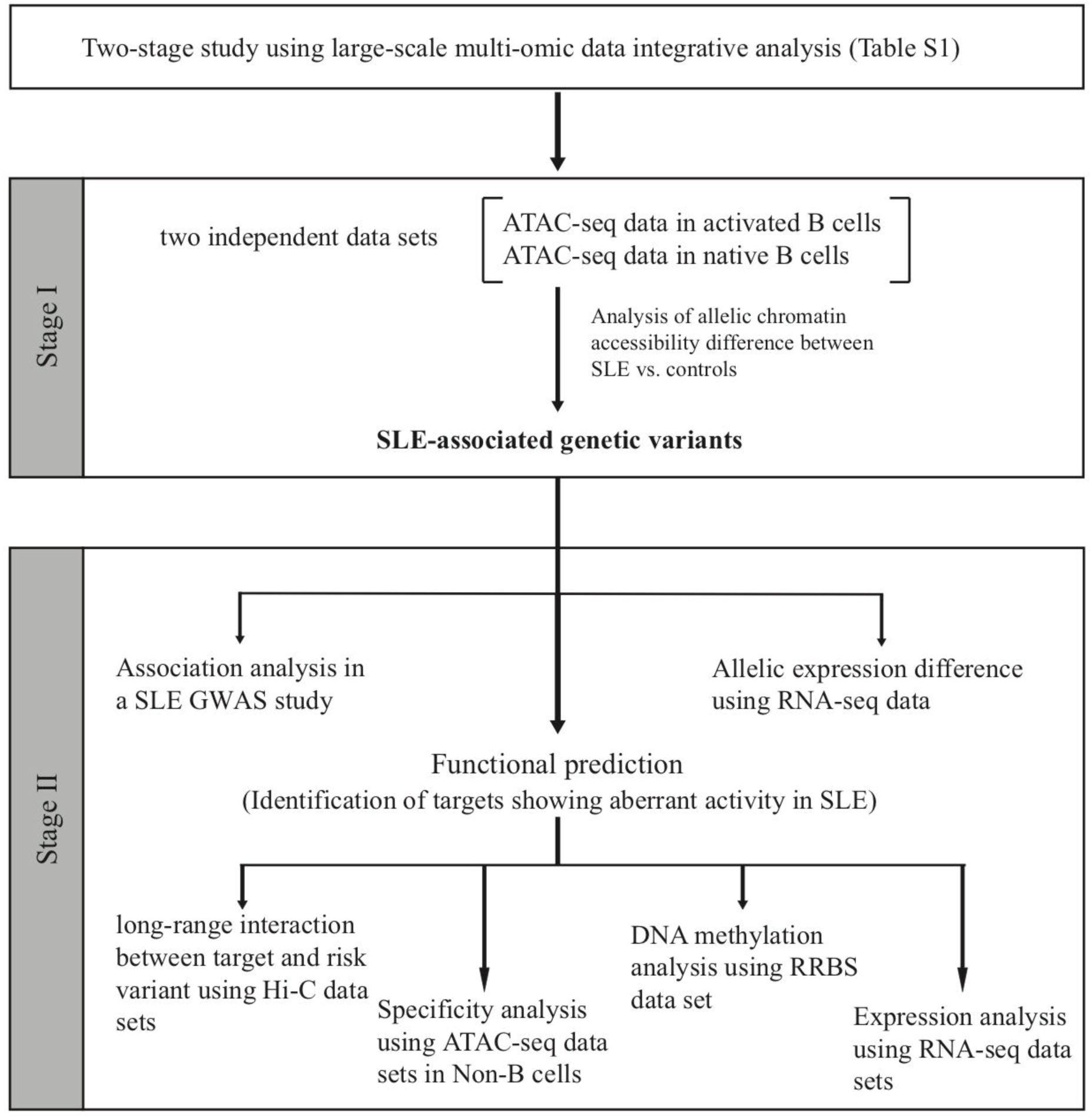
Schematic of the study design. On the basis of the functional genomic data feature, a two-stage study was designed. Summary of data sets are available in Table S1 and S2.

**Figure 2.**
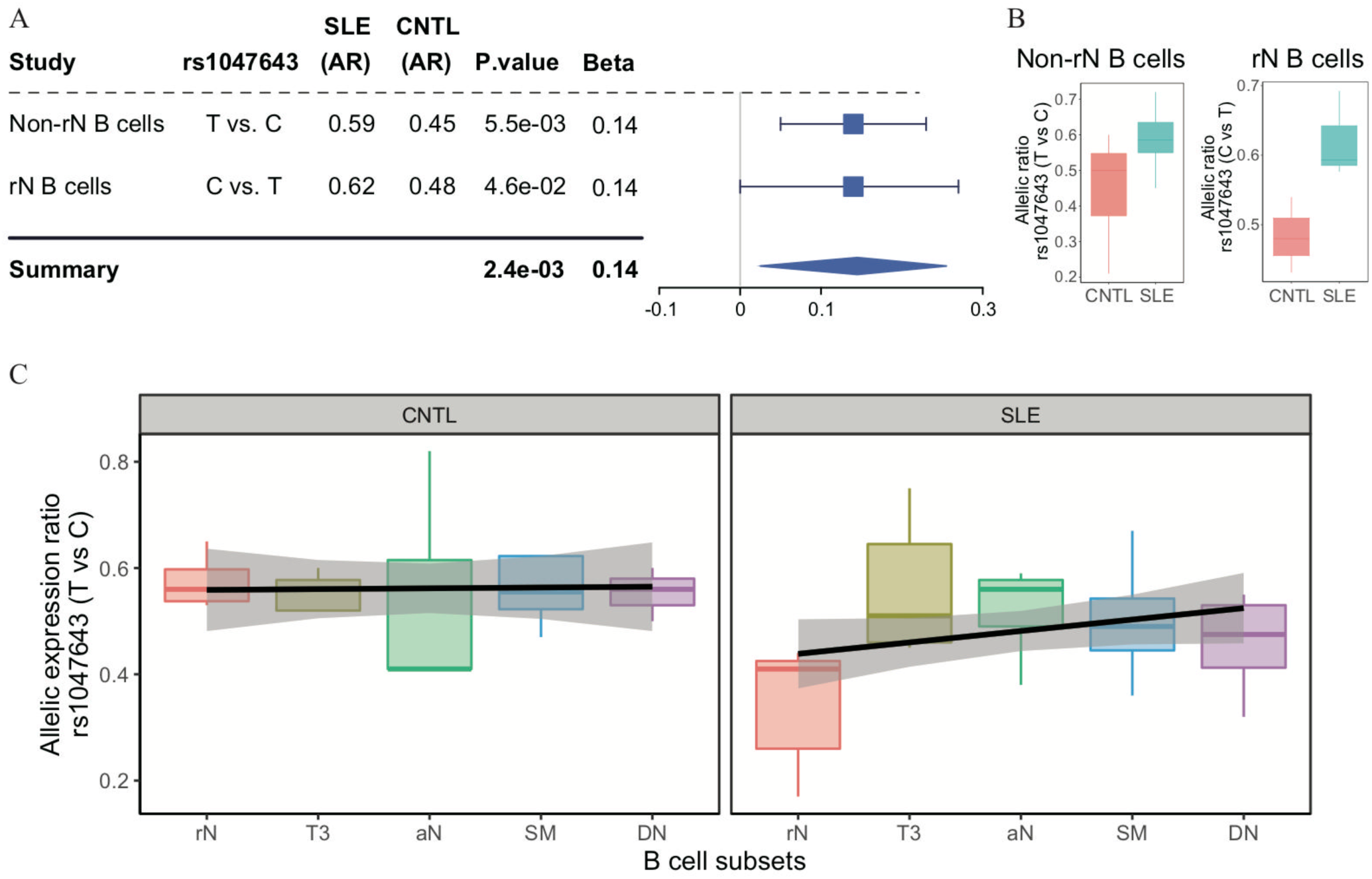
Change of allelic chromatin accessibility and expression in B cell subtypes from SLE patients and controls. (A) Forest plot showing AI of allelic chromatin state of SNP rs1047643 in both resting naive (rN) and activated (Non-rN) B cells in patients of SLE compared with healthy controls. The plot in the right panel displays the 95% of confidence interval of beta-value. (B-C) Boxplots showing allelic expression of SNP rs1047643 in both rN and activated B cells in patients with SLE as compared with healthy individuals. All raw data are available in Figure 2—source data 1.

Because the rs1047643 is located in the first exon of *FDFT1* gene (Figure 3E), it enables us to test the functionality of this variant at the transcriptional level. Analyzing RNA-seq data (Accession ID: GSE118254), we determined the AI of RNA transcripts for the rs1047643. In line with results shown above, we observed the dynamic AI pattern on the transcriptional level for the rs1047643 (Figure 2B). Meanwhile, this dynamic allelic expression pattern is specific during B cell development with SLE (Figure 2C).

**Figure 3.**
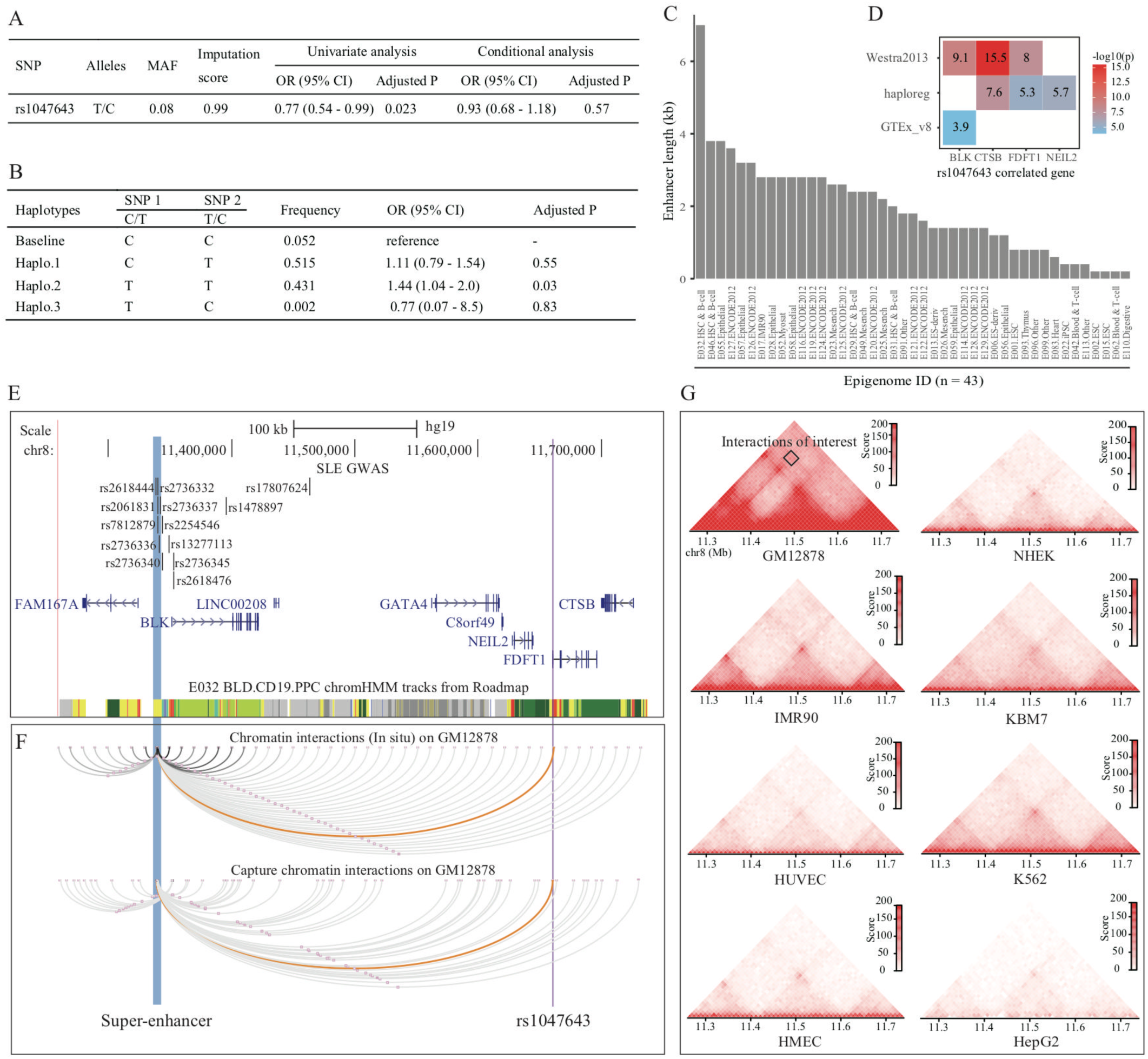
Association analysis and functional prediction of SNP rs1047643. (A) Association results for the SNP rs1047643 with SLE risk in single marker analyses. MAF, minor allele frequency; OR, odds ratio; CI, confidence interval. (B) Haplotype analyses of the two SNPs (SNP1: GWAS indexed SNP rs17807624; SNP2: rs1047643) in relation to SLE risk. Baseline (the reference haplotype) represents the alleles associated with a reduced risk in two SNPs. (C) Barplot showing the genomic length of chromHMM-annotated enhancer state on the super-enhancer region (blue highlighted in 3C) in 43 epigenomes. (D) Plot shows the eQTL result of SNP rs1047643 in whole blood or B cells from three databases (shown in y-axis). (E) Genomic annotations of the SNP rs1047643. The three tracks show locations of 13 GWAS index SNP, gene annotation and 15-state chromatin segments in CD19+ B cells at 8p23 locus, respectively. Vertical blue and purple lines, represents the location of super-enhancer and SNP rs1047643, respectively. (F) Long-range interaction between a super-enhancer and SNP rs1047643. The two tracks show chromatin interactions from two independent studies using whole-genome Hi-C and capture Hi-C technologies, respectively. Orange curves show the interactions between the super-enhancer and the SNP rs1047643. (G) Heatmaps showing the 3D DNA interactions at 8p23.1 locus in eight cell lines. The rectangle represents interactions between the super-enhancer and the SNP rs1047643. All raw data are available in Figure 3—source data 1.

### Association with SLE risk in American Hispanic populations

Because SNP rs1047643 has not been reported to be associated with the susceptibility of SLE and other autoimmune diseases, we next tested the association using a dataset from an SLE GWAS case-control study. Employing the univariate analysis for SNP rs1047643 in samples from Hispanic populations, we identified an association of the rs1047643 with SLE risk at statistical significance of adjusted *P* = 0.02 (Figure 3A), albeit not reaching the significance after adjustment for 12 GWAS index SNPs (the top track in Figure 3E, where one SNP rs2736336 is excluded due to its multivariate alleles). Of the 12 index SNPs, indeed, one index SNP rs17807624 with the statistical significance with *P* < 1.5 × 10^-3^ using the univariate analysis, is the top signal to which the SNP rs1047643 is conditional. Thus, we performed haplotype analyses on these two SNPs (index SNP rs17807624 and rs1047643, Figure 3B). Compared with the reference haplotype, which carries the alleles associated with a reduced risk in two SNPs, haplotype 2, which carries the risk-associated alleles, showed a significant association (adjusted *P* = 0.03).

### Functional annotation

An analysis of eQTL data derived from three independent cohorts indicated both proximal (< 200 kb) and distal (> 200 kb) regulatory potential for the SNP rs1047643 in normal B or blood cells (Figure 3D). Interestingly, besides correlated with three adjacent genes (*FDFT1*, *CTSB* and *NEIL2*), the rs1047643 is also an eQTL linked with an upstream *BLK* gene in a distance of ∼300 kb, a result that is detected in two independent data sets. An analysis of RNA-seq data from two independent studies (Accession ID: GSE118254 and GSE92387, Table S1) consistently showed that expression patterns for two representative genes (*BLK* and *FDFT1*) are gradually increased in a developmental process from naive to memory B cells, in particular, the double negative memory B cell subset in patients with SLE, the pattern that is not observed in controls (Figure S1 and S2).

By searching for enhancers and other regulatory elements across 8p23 locus from a dataset of the 127 epigenomes from Roadmap, we identified a SE with a length of 7kb in the upstream of *BLK* gene in CD19+ B cells (Epigenome ID: E032, Figure 3E). An analysis of annotated enhancer elements across the 127 epigenomes showed 43 (33.9%) epigenomes had enhancers at this SE region. Comparative analysis of the enhancer length at this SE region on the 43 epigenomes further showed that this SE is specific in CD19+ B cells (Epigenome ID: E032, Figure 3C).

Analyzing Hi-C data sets from two independent studies in GM12878 cells, we observed a DNA looping between the SNP rs1047643 within FDFT1 and the SE region (Figure 3F). More importantly, in GM12878 B-lymphoblastic cells, this SE region has a wealth of long-range interactions with adjacent genes (e.g., BLK) and functional elements. In contrast, in another seven cells (Figure 3G), as well as in normal T cells (Figure S3) and nine selected tissues (Figure S4), these interactions are either much weaker or completely absent. These results indicate that the physical interaction between SNP rs1047643 and SE region, and many interactions with this SE, are specific to B-lymphocytes.

### Specificity in B cells

We then hypothesized that the SE region may show aberrant activity in B cells from SLE patients. To test this hypothesis, we conducted quantitative analysis on the same ATAT-seq data (Accession ID: GSE118253 and GSE71338) used in stage I (see Methods in detail). Comparison of SE activity in a quantitative manner between SLE patients and controls indicated that the SE activity is gradually increased through B cell development in SLE patients (Figure 4A-B), with a hyper-activity being observed in double negative (DN) B cells in patients, relative to controls (Figure 4B-C).

**Figure 4.**
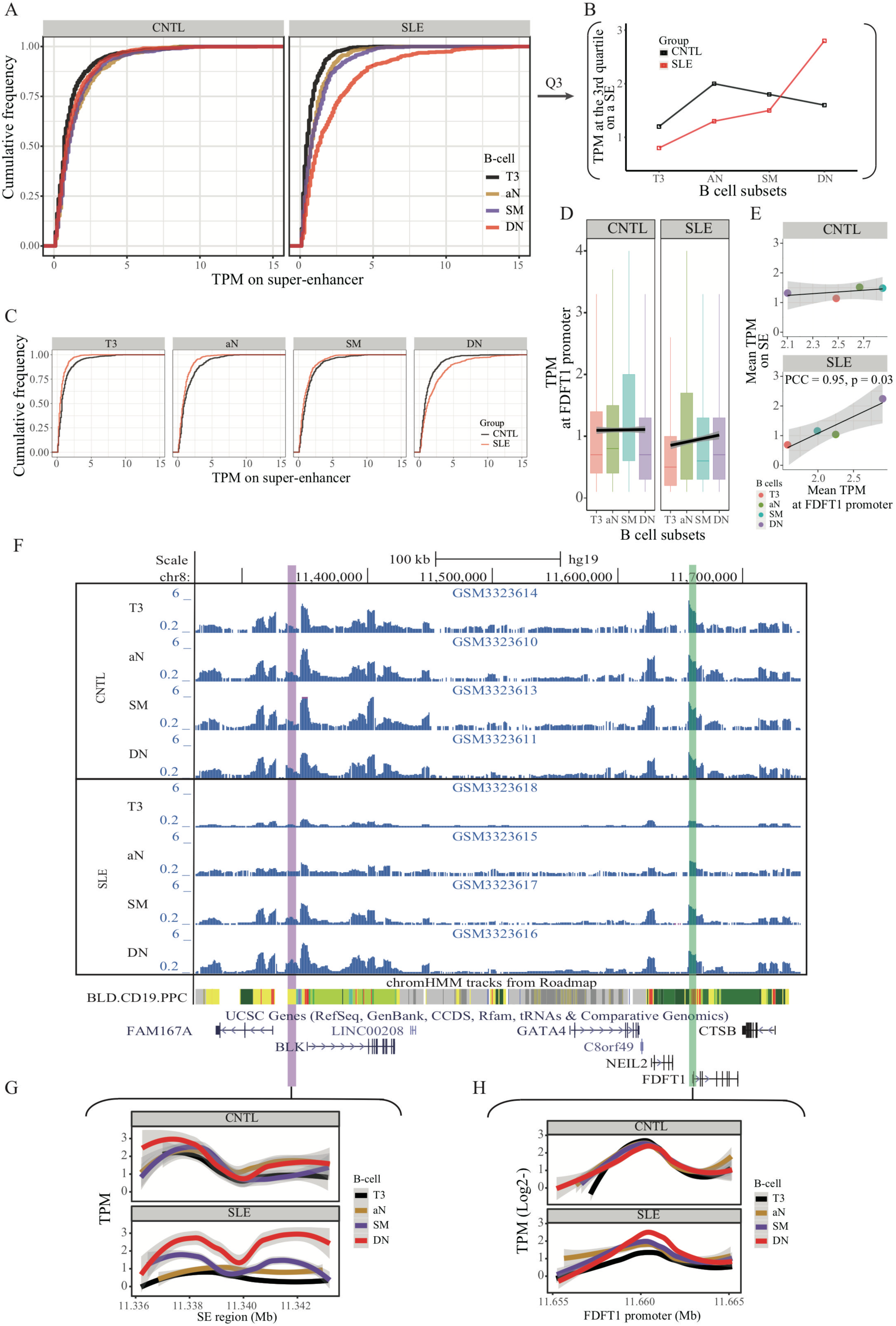
Aberration of super-enhancer and *FDFT1* promoter region in B cell subtypes from SLE patients. (A) Empirical cumulative distribution of TPM values per 50-bp window across the 7kb SE region in B cell subsets for disease and control groups. (B) Plots showing the TPM values at the third quartile (Q3) across B cell subtypes as a comparison between SLE and controls. (C) Empirical cumulative distribution of TPM values on the SE region (same as shown in A) in a comparison between two groups across four B cell subtypes. (D) Boxplots showing the TPM values per 50-bp window at the *FDFT1* promoter region in B cell subtypes for SLE and controls. The black lines and grey areas represent the linear regression results towards the B cell development from T3 to DN stages, and 95% of CI. (E) Plots showing the correlation between super-enhancer and *FDFT1* promoter regions based on mean TPM values with respect to B cell subtypes in SLE and controls. (F) Wiggle plot showing the enrichment of open chromatin states at 8p23.1 locus in B cell subtypes for two individuals (a healthy individual at upper panel, and a patient with SLE at lower panel). Purple and green vertical lines represent the locations for super-enhancer and *FDFT1* promoter, respectively. Quantitative comparison of chromatin accessibility states in SE (G) and *FDFT1* promoter regions (H) with respect to B cell subtypes. All raw data are available in Figure 4—source data 1.

Similarly, the rs1047643-containing promoter activity also shows up-regulation towards B cell development in SLE patients (Figure 4D-E). In a comparison of B cell development on activities of SE and FDFT1 promoter regions in two individuals, the chromatin accessibility on both regions in an individual with SLE is increased during B cell development, but remains relatively unchanged in the healthy individual (Figure 4F-H).

We also quantitatively compared open chromatin states of SE and FDFT1 promoter regions in resting naive B cells (Accession ID: GSE71338). Concordant with the results from active B cell subsets, the open chromatin states on both regions are low in non-active B cells from SLE patients, relative to healthy controls (Figure S5).

We further conducted quantitative analyses on ATAC-seq data from another two independent studies in two immune cell types, T cells and neutrophils (Accession ID: GSE139359 and GSE110017, Table S1). The results showed that there was no marked enrichment of ATAC-seq reads on both the SE and FDFT1 promoter regions in these two immune cell types for both SLE and controls (Figure S6). Collectively, these results suggest a B cell specific, rs1047643-interacting SE whose activity is aberrant in SLE B cell development.

### Hypomethylation in SLE B cells

We further analyzed DNA methylation in the SE region using RRBS data in B cell development in a comparison between SLE and controls (Accession ID: GSE118255, Table S1). Our results show that DNA methylation levels on the SE region are gradually decreased in the developmental process from resting native (rN) to memory B cells in patients with SLE (Figure 5A). In contrast, there is no such obvious change of DNA methylation pattern in the control group. A correlation analysis also showed a marked negative correlation between open chromatin states (TPM values, also presented on Figure 4E) and DNA methylation levels at the SE region in the SLE group, relative to the healthy controls (Figure 5B). Together, these results reinforce the aberrant activity of SE in developmental process of B-lymphocytes in patients with SLE.

**Figure 5.**
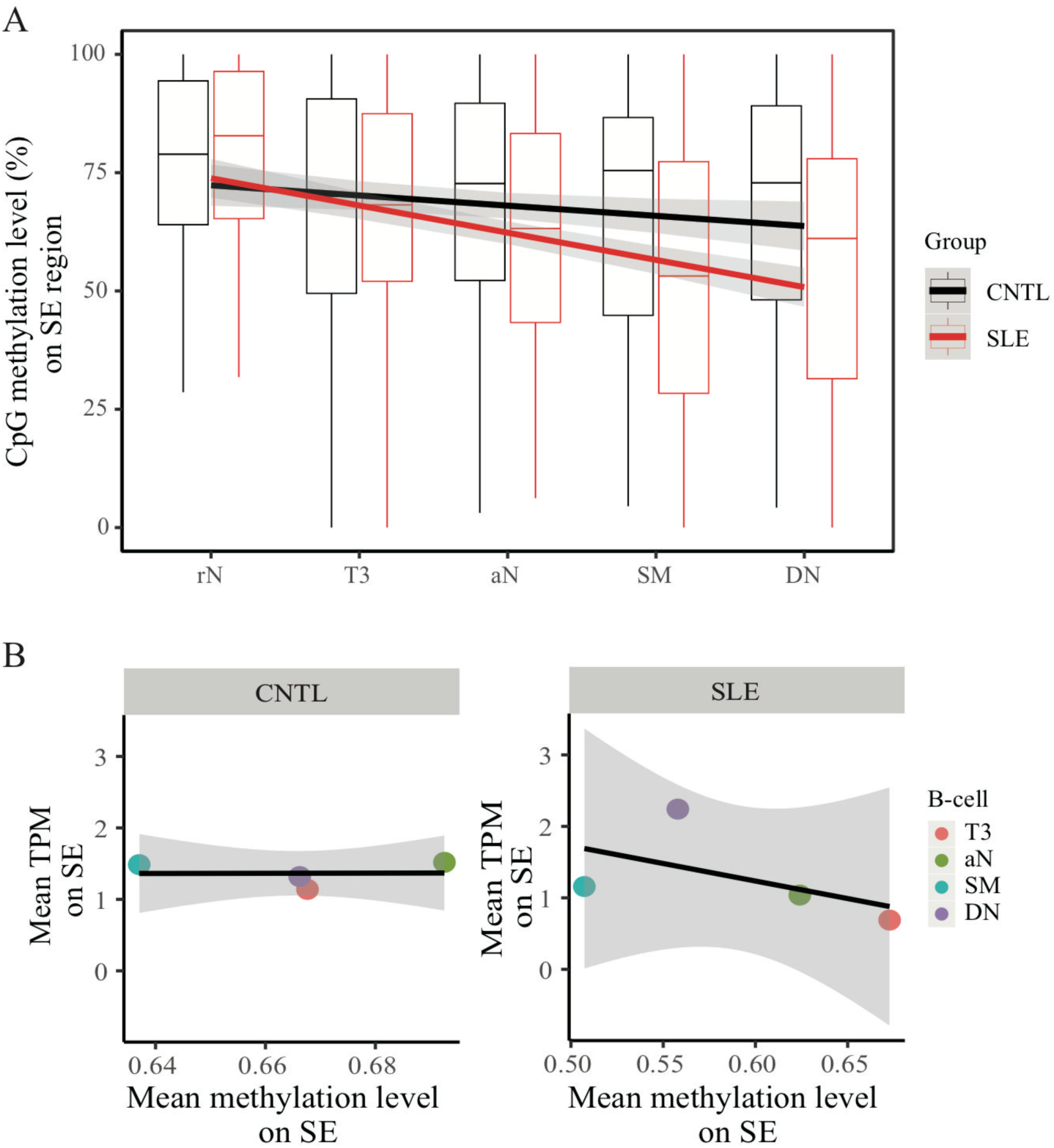
Hypomethylation in super-enhancer region in B cell subtypes from SLE patients. (A) Boxplots showing the CpG methylation levels per 50-bp window in 7kb SE region in B cell subtypes for SLE and control groups. The black and red lines represent the linear regression results towards the B cell development from rN to DN stages for SLE and controls, respectively. (B) Plots showing the correlation between TPM values (y-axis) and DNA methylation levels (x-axis) averaged over each B cell type in SLE and controls. All raw data are available in Figure 5—source data 1.

### STAT3 binding on both super-enhancer and rs1047643-residing regions

TF-motif enrichment and binding analysis using the ENCODE TF ChIP-seq dataset (v3) predicted that STAT3 may bind to both the SNP rs1047643-containing promoter and SE regions (data not shown). To validate the finding, we designed two pairs of primers (SE5 and SE3, Figure 6A) to determine the STAT3 binding on SE region and its contribution to the SE activity using STAT3, H3K4me1 and H3K27ac ChIP-qPCR assays in GM11997 cells. Under normal culture conditions, we validated that pSTAT3, H3K4me1 and H3K27ac modifications are remarkably enriched on the SE region in B-lymphoblastic cells, relative to IgG mock controls (Figure 6A, 6E and 6F). We then conducted both the inhibition and activation of STAT3 DNA binding activity using two small molecules. In B-lymphoblastic cells challenged with S3I-201, a STAT3 DNA binding inhibitor, both the DNA binding of STAT3 on SE region and the SE activity are significantly reduced (Figure 6A), relative to control. In GM11997 cells treated with ML115, a selective activator of STAT3 (36), both the STAT3 DNA binding capability on SE region and the SE activity are significantly increased (Figure 6E), relative to controls. These results together demonstrate that STAT3 directly modulates the SE activity.

**Figure 6.**
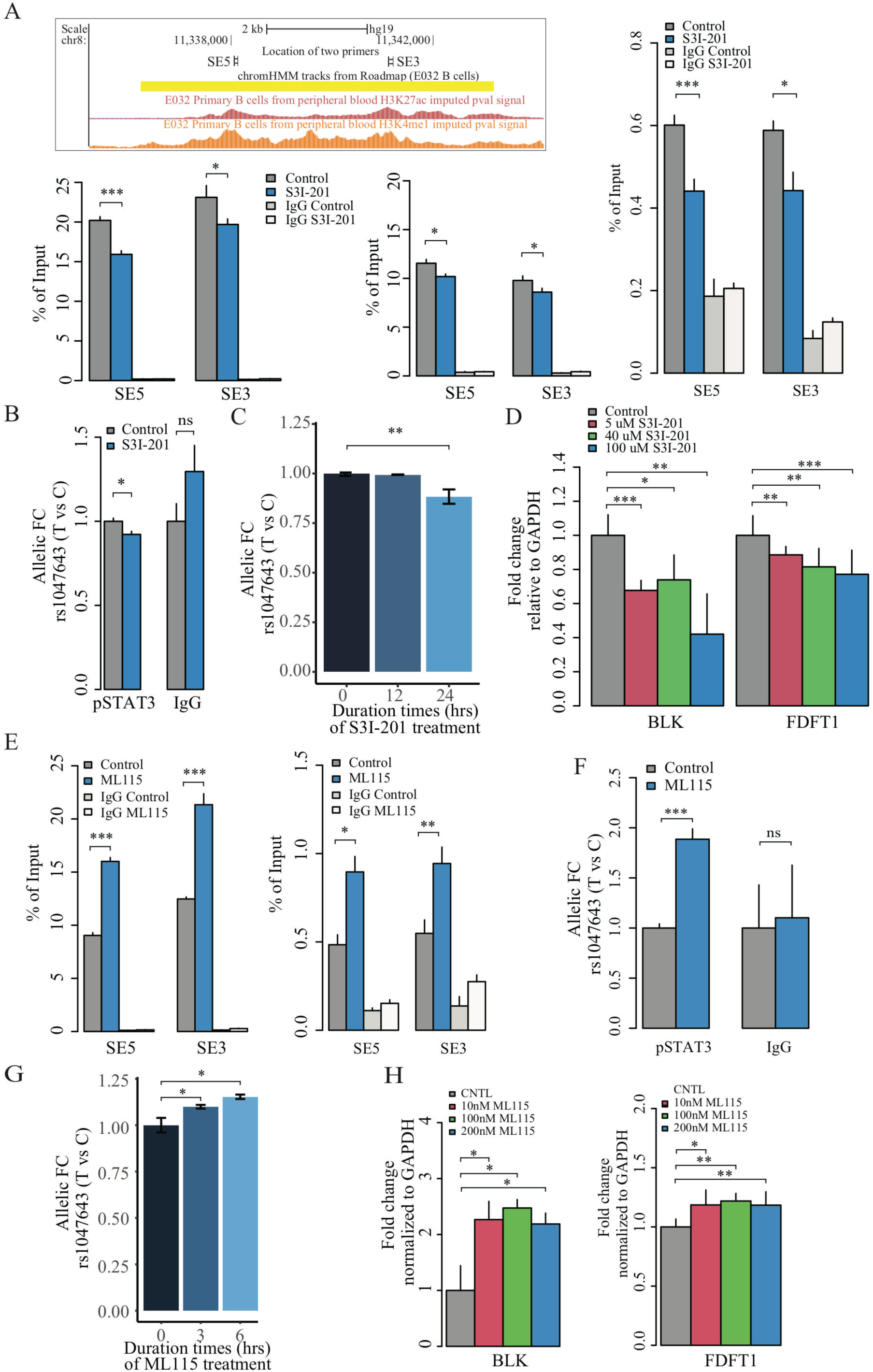
Contribution of STAT3 modulates the enhancer activity and SNP-residing locus in cultured GM11997 cells. (A) ChIP-qPCR for H3K27ac (left lower panel), H3K4me1 (middle lower panel) and pSTAT3 (right) at 8p23 super-enhancer region following 40 μM S3I-201 treatment for 24h. Left upper panel: UCSC genome browser showing the location of two pairs of qPCR primers (SE5 and SE3) on the SE region (yellow). Two tracks shown below are the enrichment of H3K27ac and H3K4me1 across the SE region. (B) Allelic ChIP-qPCR for pSTAT3 binding and (C) allelic RT-qPCR on SNP rs1047643 (T vs C alleles) following 40 μM S3I-201 treatment for 24h. (D) RT-qPCR with RNA from B-lymphoblastic cells that have been challenged with S3I-201 for 24h as indicated. The fold changes for the rs1047643-associated *BLK* and *FDFT1* genes in response to different concentrations of S3I-201 compared to vehicle (0.1% DMSO) as control, which was set as 1 in all cases, are presented. (E) ChIP-qPCR for H3K27ac (left), and pSTAT3 (right) at 8p23 super-enhancer region following 100 nM ML115 treatment for 6h. (F) Allelic ChIP-qPCR for pSTAT3 binding and (G) allelic RT-qPCR on SNP rs1047643 (T vs C alleles) following 100 nM ML115 treatment for 6h. (H) RT-qPCR with RNA from B-lymphoblastic cells that have been challenged with ML115 for 6h as indicated. The fold changes for the rs1047643-associated *BLK* and *FDFT1* genes in response to different concentrations of ML115 compared to vehicle (0.1% DMSO) as control, which was set as 1 in all cases, are presented. NS, not significance; *, *P* < 0.05; **, *P* < 0.01; ***, *P* < 0.005.

We next tested whether the STAT3 might also regulate the rs1047643-residing regions. Using allelic qPCR assay, we confirmed that genomic DNA in the GM11997 cells carries a heterozygous variant for the SNP rs1047643 (Figure S7), enabling the AI analysis in this cell model. In GM11997 cells treated with the STAT3 inhibitor S3I-201, STAT3 binding on the risk allele T is significantly reduced, relative to the rs1049643-C allele (Figure 6B). Concordantly, the expression level on the rs1049643-T allele is also declined after treatment with S3I-201 for 24 hours, relative to the C allele (Figure 6C). Conversely, we observed an increase of both STAT3 DNA binding and expression at the rs1049643-T allele in cells stimulated with the STAT3 activator ML115 (Figure 6F-G). These results collectively suggest that the risk rs1049643-T allele is preferentially bound by STAT3 in B cells.

We also determined RNA expression of BLK and FDFT1, two representative genes that correlate with the risk rs1047643. The expression levels of both genes are decreased with the treatment of S3I-201 (Figure 6D), and up-regulated with the STAT3 activator ML115 (Figure 6H). These results suggest the STAT3-binding risk allele T is associated with increased expression of *BLK* and *FDFT1*.

## Discussion

In the present study, by integrating a variety of functional genomic data, we performed AI analysis to uncover novel functional promising variants and their regulatory targets in association with SLE. Of note, the diversity of genomic data types from this comprehensive data collection for autoimmune diseases allowed us to develop an approach not used before for accessing the role of variants in SLE disease activity.

One of the most significant findings is the identification of a novel risk variant rs1047643. The association study shows that the rs1049643-T is a risk allele for SLE. Our AI analyses indicate that the rs1049643-T allele resides in more open chromatin state and has higher expression in SLE memory B cell subsets, relative to the C allele. Functional study further provides evidence that the rs1049643-T allele is preferentially bound by STAT3. The SNP rs1047643 is also an eQTL linked with both proximal and distal genes, including *BLK*, the gene that plays a critical role in B lymphocyte development (37). These results demonstrate that this novel SLE-associated risk rs1047643 whose functionality is mediated by STAT3, may play a role in allele-specific control of adjacent genes at 8p23 locus in B cells. Despite no report for association with other autoimmune diseases, this SNP has been associated with multiple myeloma (38) and follicular lymphoma (39), two malignant diseases whose pathogenesis is partially associated with the dysfunction of B cells. Specifically, hyperactive STAT3 has been reported to be associated with poor survival in both diseases (40, 41). Therefore, our findings may provide a clue for genetic and mechanical studies on those B cell associated diseases.

Another intriguing finding in this study is the identification of an aberrant activity of a SE in lupus B cell subsets, particularly the hyperactivity in memory B cells. In contrast, there is no enhancer activity in other immune cells (T cells and neutrophils analyzed in this study) in patients with SLE. We also demonstrate that the aberrant activity of the SE can be mediated by STAT3. Some studies have consistently reported a critical role of STAT3 in the B cell maturation, differentiation, as well as the autoimmunity (42, 43). These reports further support the significance of STAT3-mediated SE aberration in B cells with SLE.

Several studies have highlighted the 8p23 locus as a major SLE susceptibility region (44). Our study further expands the significance at this locus. We speculate that the 8p23 locus may play functional roles in B cell development in both genetic and epigenetic fashions. Besides the SNP rs1047643 discovered in the present study, there are 13 SLE-associated GWAS leading SNPs reported in this locus. Of 13 SNPs, six SNPs (Figure 3E) directly sit in the SE region, suggesting these risk variants may play roles in a genetic interaction way. For example, our study and others together suggest that there are a few cis-eQTLs linked with transcriptional levels of BLK (11, 44). Epigenetically, the SLE-associated SE has physical interactions with adjacent genes, including *BLK* and *FDFT1*, and the risk rs1047643-residing region. This indicates a potentially complex role of the variant rs1047643 for broad regulation by physically contacting the SE. Thus, our data provide new insights into the molecular mechanisms by merging genetic susceptibility with epigenetic impacts on gene expression for autoimmune diseases.

The *FDFT1* is a gene encoding for squalene synthase, the enzyme that catalyzes the early step in the cholesterol biosynthetic pathway (45). Previous studies have shown dyslipidemia, with elevations in total cholesterol, low-density lipoprotein, triglyceride levels in patients with lupus (46), especially in the active disease. Our multi-omics data indicate that the SNP rs1047643-linked FDFT1 is aberrantly activated in B cell development in SLE patients, thereby providing an insight into the genetic implication of lipid metabolism for autoimmune diseases.

The limitations of this study include, due to the presence of six SLE GWAS tagging SNPs in SE region, we are unclear how they genetically influence the SE activity during B cell development. Second, it remains unclear how the AI pattern occurs in naive B cells with lupus. The C allele shows more open chromatin state in SLE naive B cells, this can’t be explained by STAT3 allelic DNA binding at the T allele. This implies that some other factors may also contribute to this dynamic AI pattern.

In conclusion, we identified a novel functional variant and B cell specific SE in association with the SLE pathogenesis, both mediated by STAT3, and influencing their gene targets. This insight into the mechanism by which manipulation of STAT3 affects the SE activity and its associated gene expression in B cells may have implications for future drug development in autoimmunity.

## Author contributions

Y.Z. conceived and designed the study, collected and analyzed the data, conducted the experiments, wrote the manuscript. D.A. contributed materials and data, and assisted in data analysis, interpreted the data and edited the manuscript. All authors read and approved the final manuscript.

## Declaration of interests

The authors declare no competing financial interests.

## Funding

This study was supported by the HudsonAlpha Institute Fund. The funders had no role in study design, data collection and analysis, decision to publish, or preparation of the manuscript.

## Acknowledgments

We thank Dr. Le Su for insightful suggestions, technical discussions and critical reading of the manuscript. This study was supported by the HudsonAlpha institutional funds. The funders had no role in study design, data collection and analysis, decision to publish or preparation of the manuscript.

## Supplementary data

**Figure 2—source data 1**

**Source files for presenting results in Figure 2.**

This zip archive contains all source data used for the quantitative analyses shown in Fig. 2.

**Figure 3—source data 1**

**Source files for presenting results in Figure 3.**

This zip archive contains all source data used for the quantitative analyses shown in Fig. 3.

**Figure 4—source data 1**

This txt file contains source data used for the quantitative analyses shown in Fig. 4.

**Figure 5—source data 1**

This txt file contains source data used for the quantitative analyses shown in Fig. 5.

---

**Figure S1.**
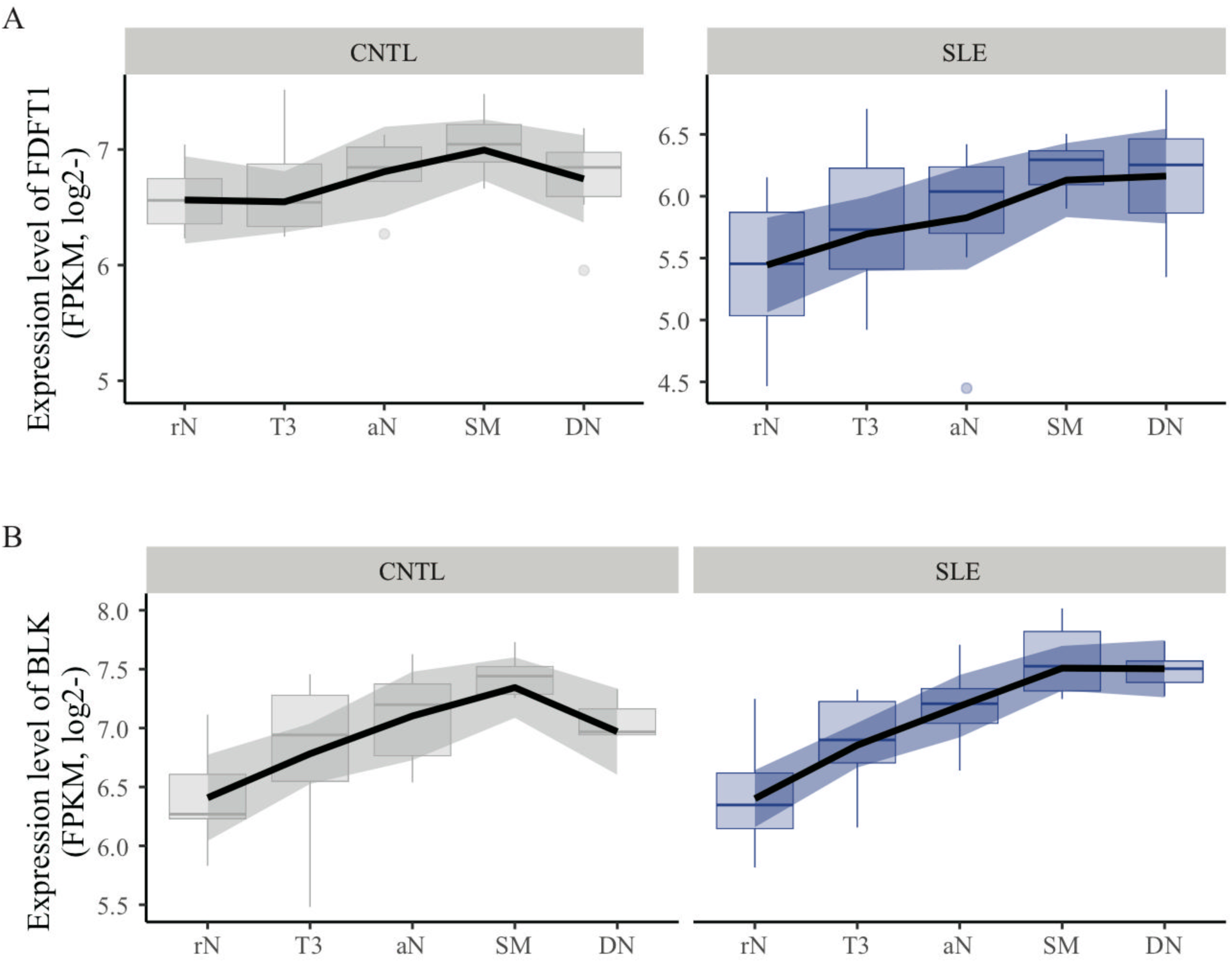
Expression pattern of FDFT1 and BLK across B cell subtypes in a comparison from a case-control study. Comparison of FDFT1 (A) and BLK (B) expression profiles in B cell subtypes from patients with SLE and healthy individuals (Accession ID: GSE118254).

**Figure S2.**
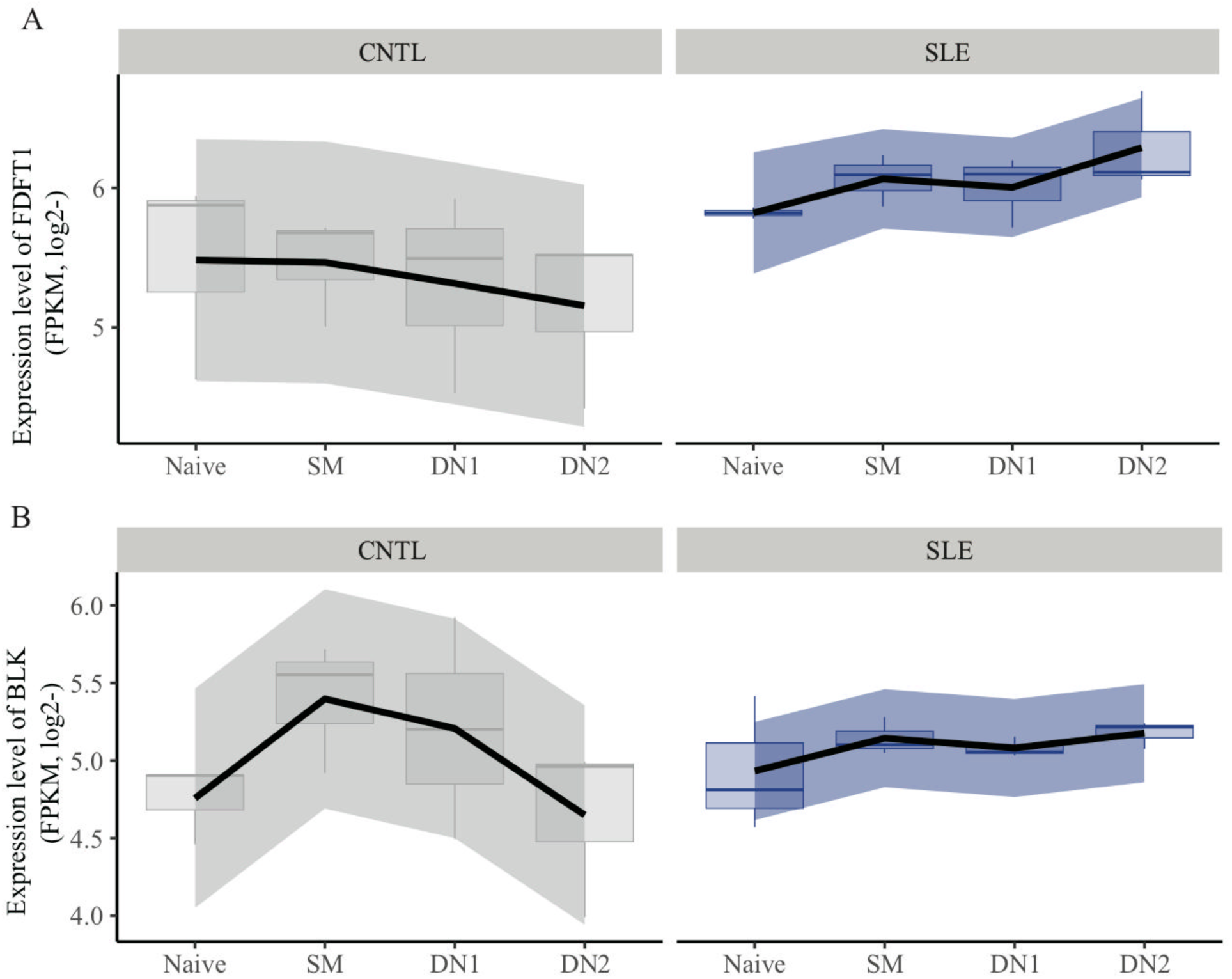
Expression pattern of FDFT1 and BLK across B cell subtypes in a comparison between patients with SLE and healthy controls. Comparison of FDFT1 (A) and BLK (B) expression profiles in B cell subtypes from a case-control study (Accession ID: GSE92387).

**Figure S3.**
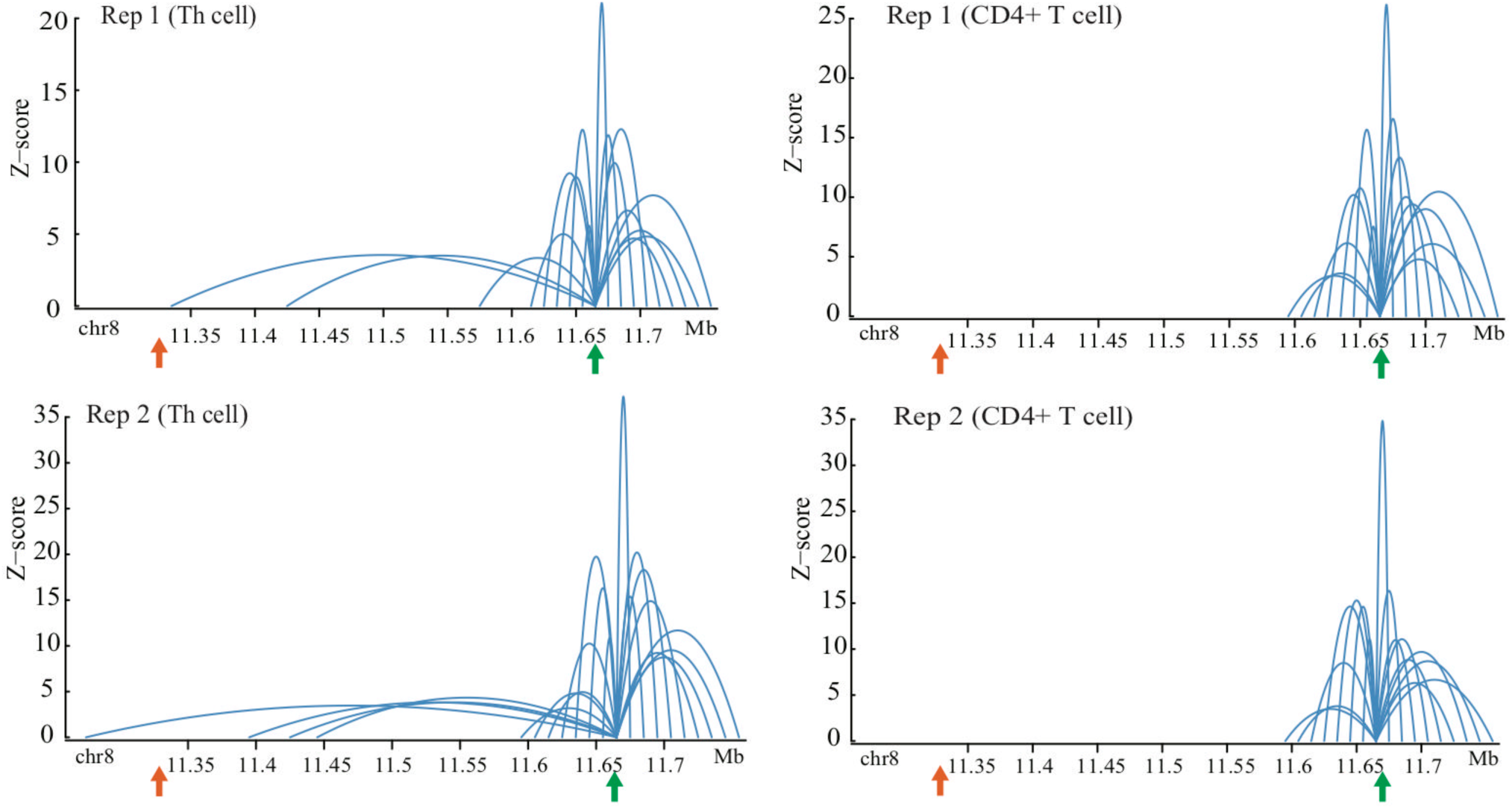
Chromatin interactions with *FDFT1* promoter region (marked in green arrow) on 8p23 locus from CHi-C data with duplicates in two types of normal T cells. Orange arrow represents the location of super-enhancer identified in this study.

**Figure S4.**
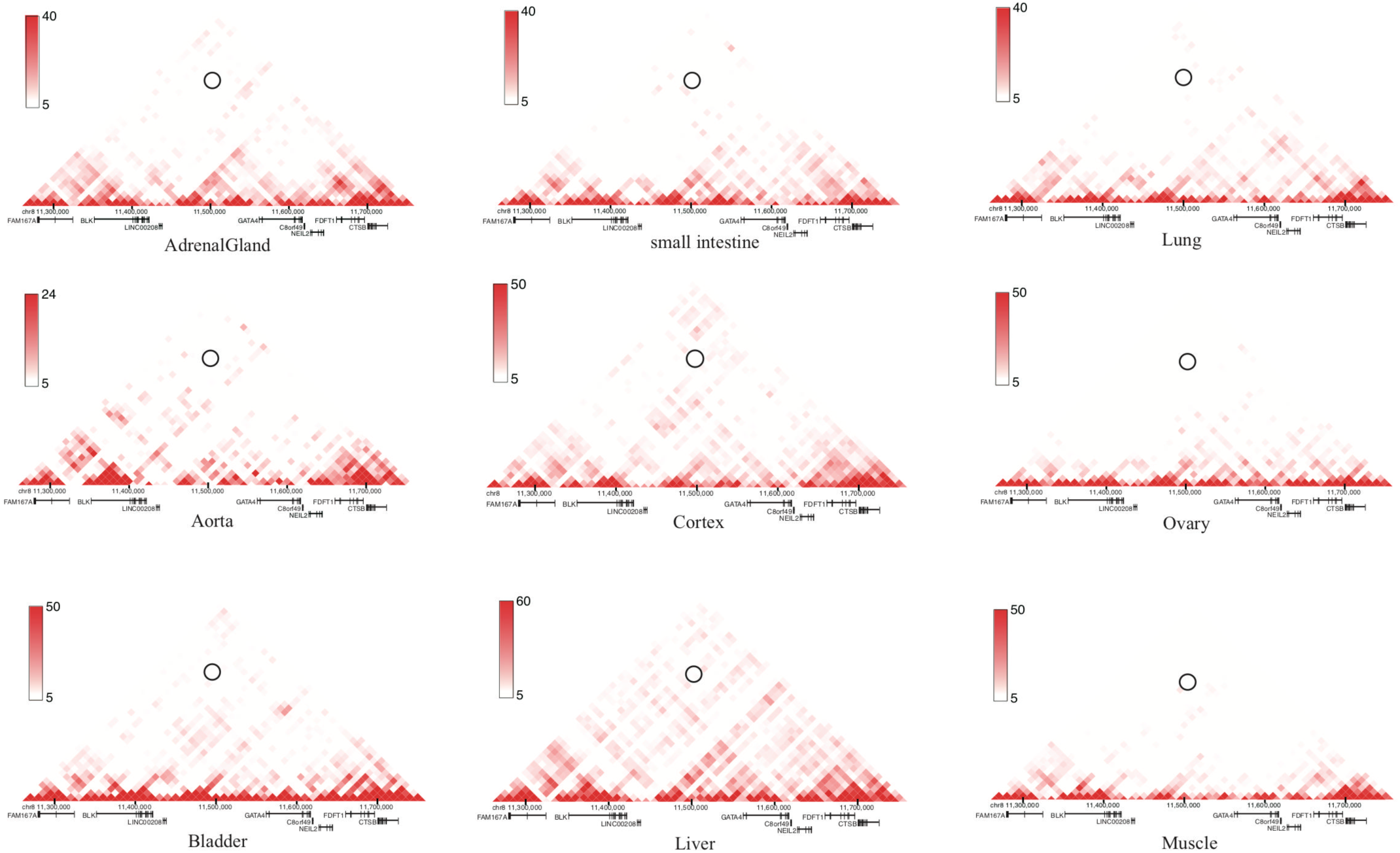
Heatmaps of Long-range chromatin interactions from Hi-C data in 8p23 locus at 10 kb (or 20 kb) resolution in a panel of human tissues from the 3D Genome Browser. The circles shown on heatmaps are the interaction score between SNP rs1047643 and SE region.

**Figure S5.**
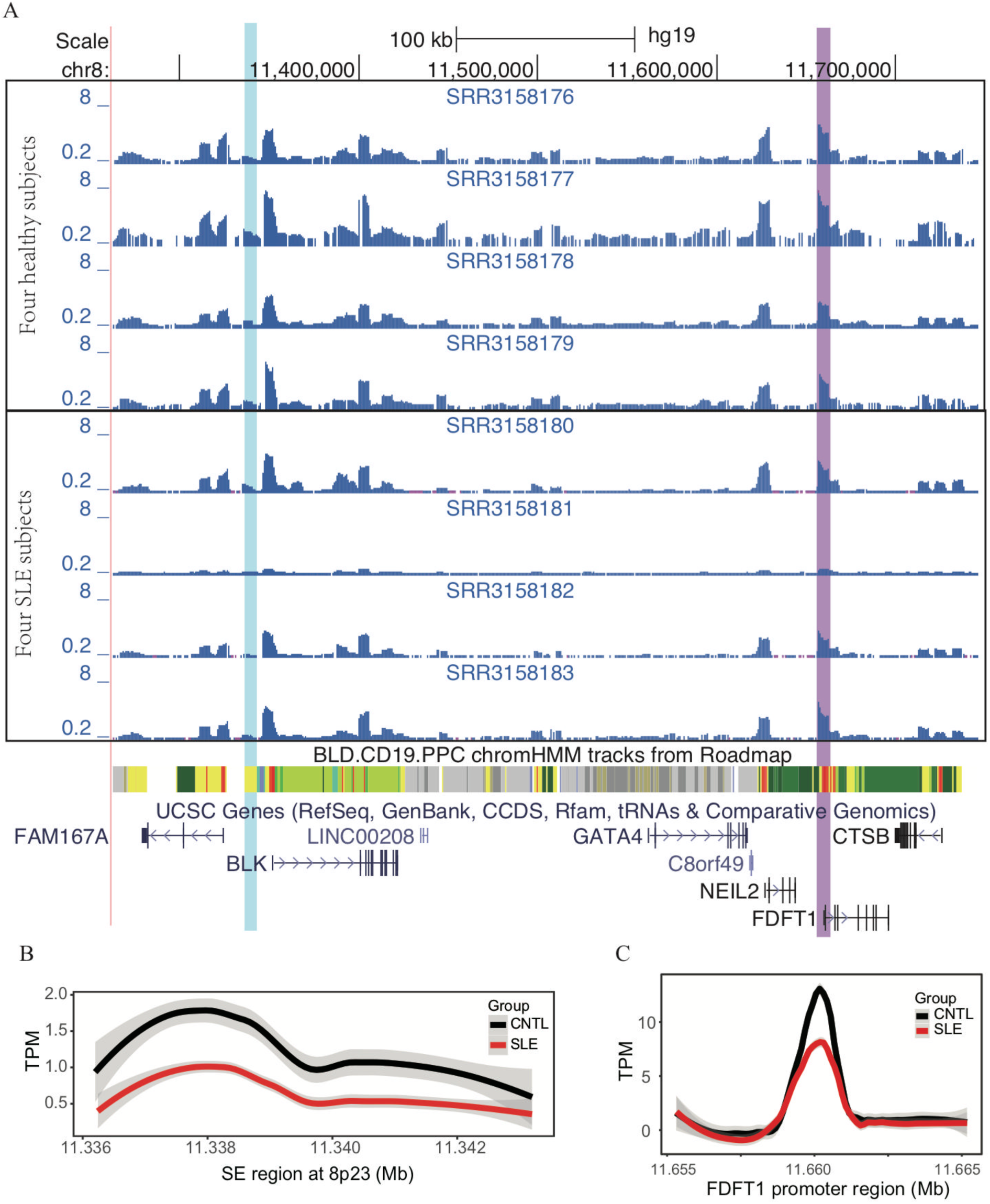
Aberration of super-enhancer in resting naive B cell subtypes from SLE patients in relation to healthy controls. (A) Wiggle plot showing the enrichment of open chromatin states at 8p23.1 locus in resting native B cells from eight individuals. Blue and purple vertical lines represent the locations of SE and *FDFT1* promoter, respectively. (B-C) Quantitative comparison of chromatin accessibility states in the SE and FDFT1 promoter regions in naive B cells in a comparison between SLE and controls.

**Figure S6.**
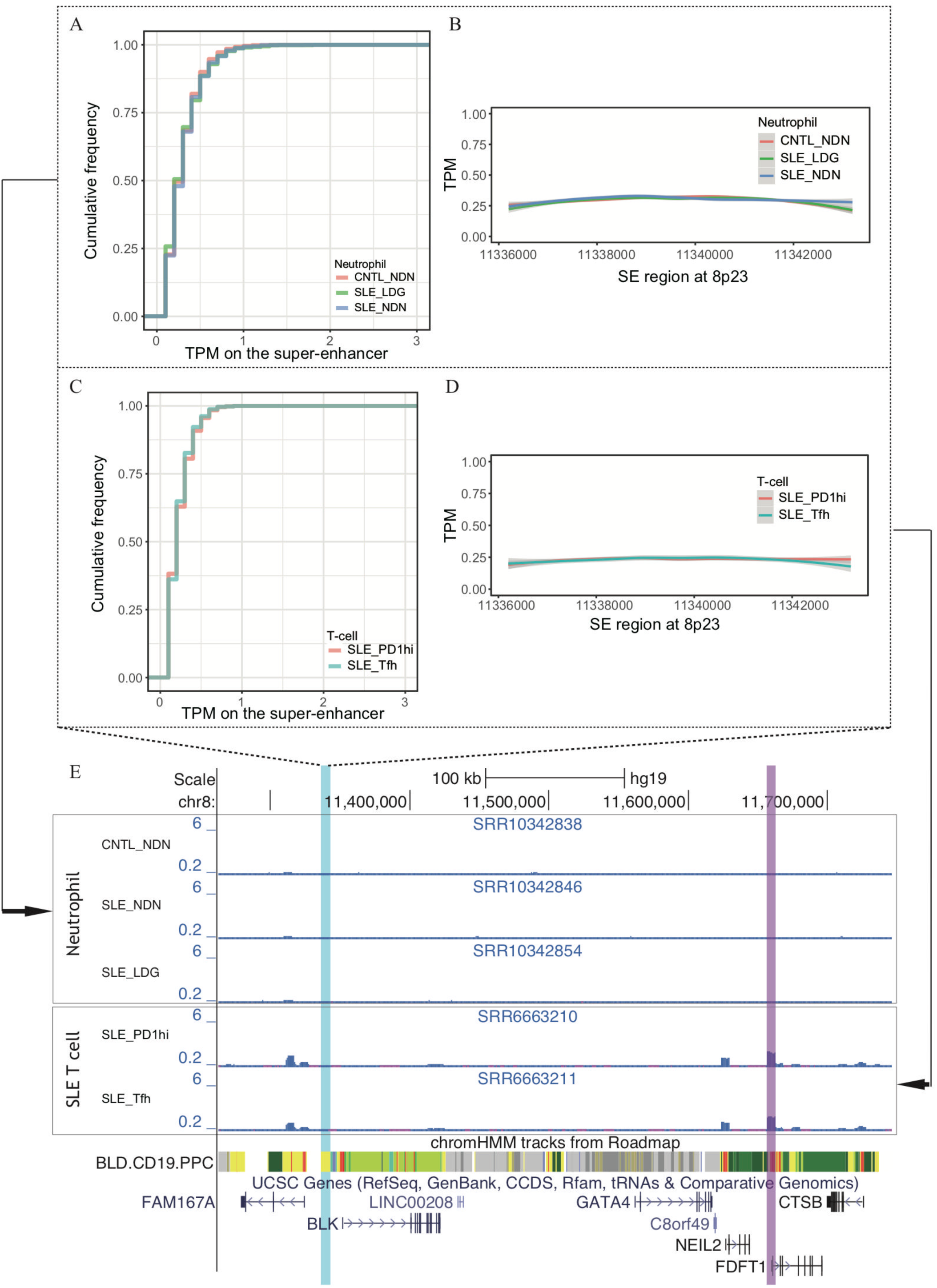
No super-enhancer activity in T and neutrophils from SLE patients and controls. (A-B) Empirical cumulative distribution of TPM values per 50-bp window and enrichment of ATAC-seq reads (TPM value) across the SE region in neutrophil cell subsets from SLE patients and controls. (C-D) Empirical cumulative distribution of TPM values per 50-bp window and enrichment of ATAC-seq reads (TPM value) across the SE region in two T cell subsets from SLE patients. (E) Wiggle plot showing the enrichment of open chromatin states at 8p23.1 locus in neutrophils and T cells. Blue and purple vertical lines represent the locations of SE and *FDFT1* promoter, respectively.

**Figure S7.**
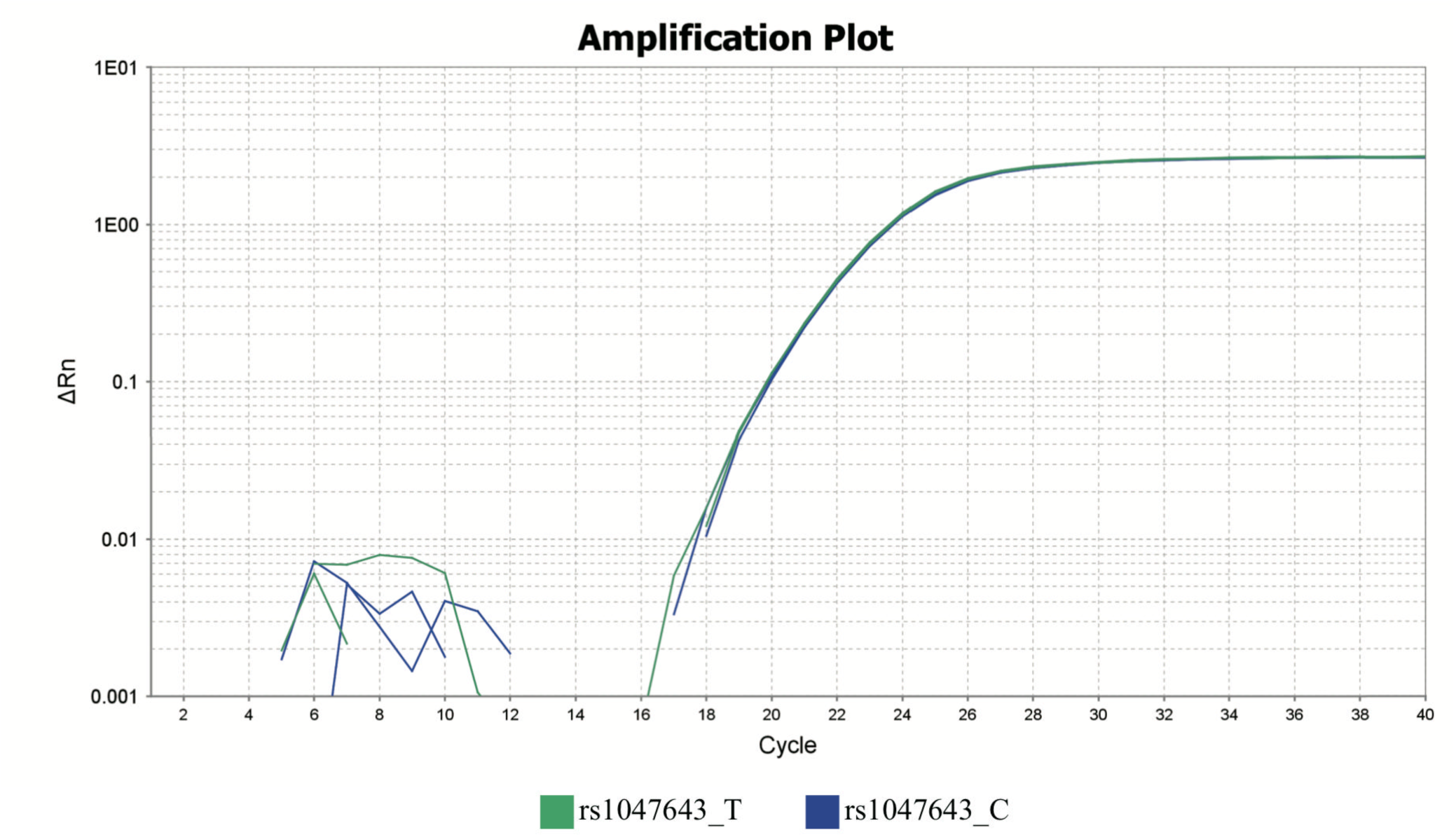
Genotyping of SNP rs1047643 in GM11997 genomic DNA using allelic qPCR analysis. Amplification plots are presented for two alleles.

**Table S1.**
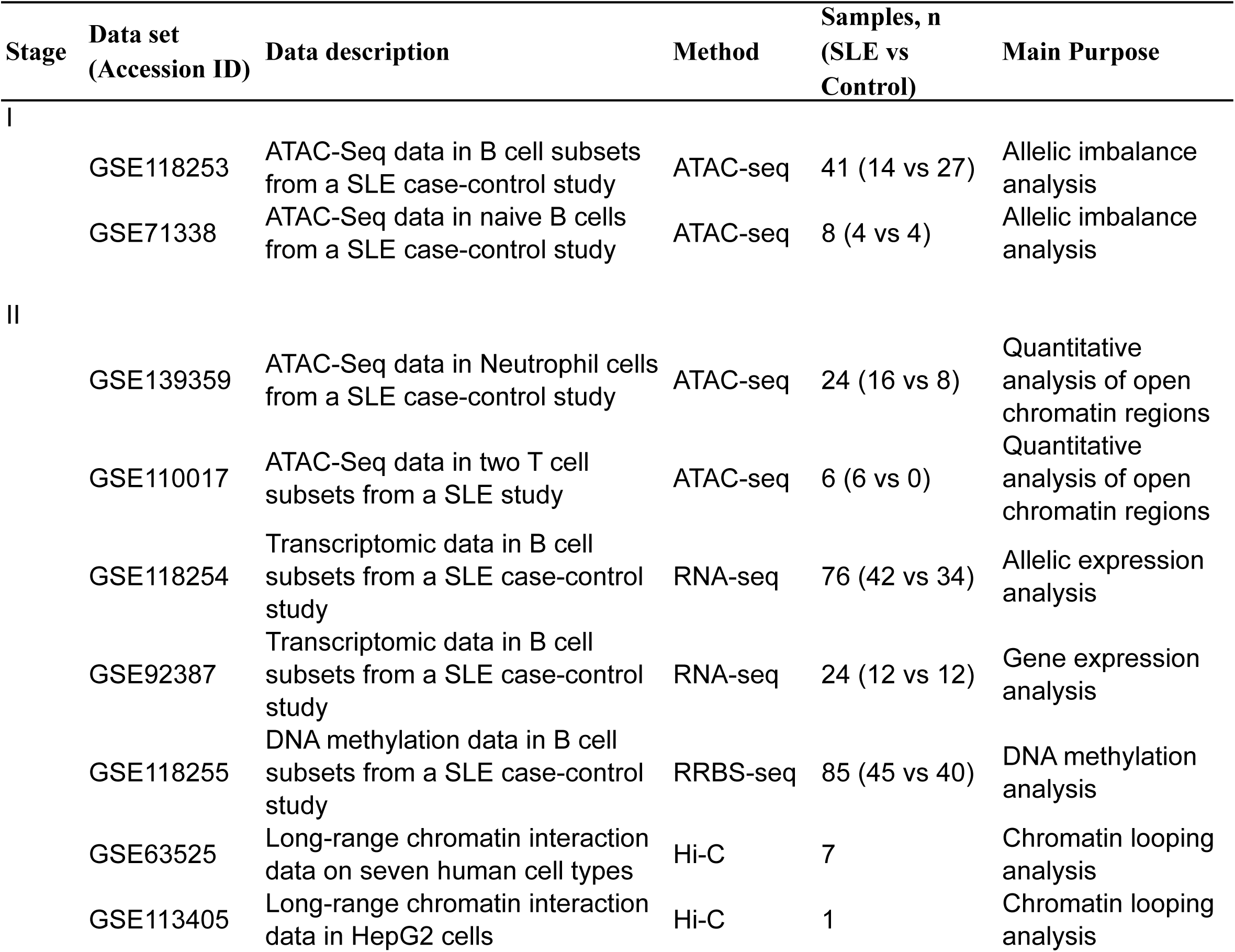
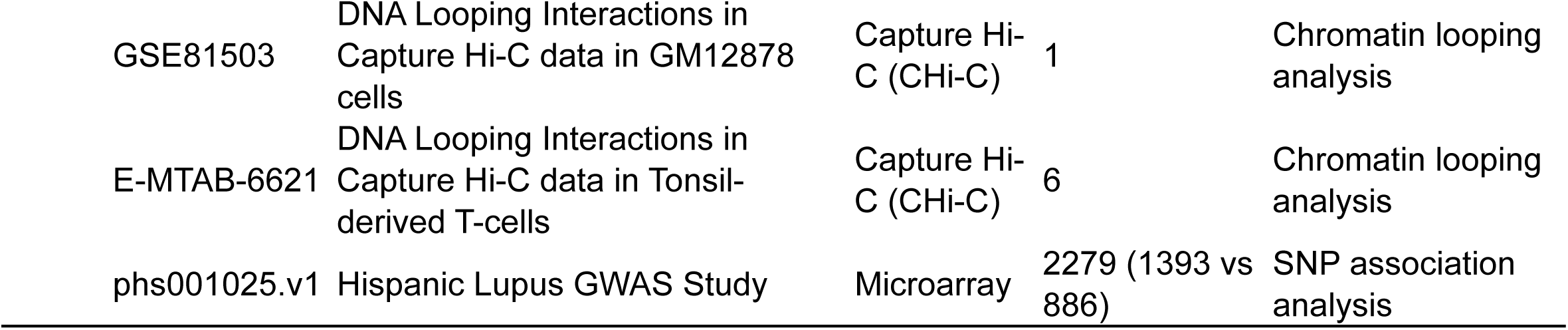
Summary of data sets used in the study. Functional genomics data sets, including ATAC-seq, RNA-seq and RRBS-seq data sets from seven SLE case-control studies (Table S2), and Hi-C data sets in multiple cell lines, and a SNP microarray data set from a lupus GWAS study.

**Table S2.** List of data sets from seven SLE case-control studies.

| **GSE ID** | **GSM ID** | **Group** | **Assay Type** | **Cell type** |
| --- | --- | --- | --- | --- |
| GSE118253 | GSM3323580 | CNTL | ATAC-seq | Activated Naive B cells (aN) |
| GSE118253 | GSM3323581 | CNTL | ATAC-seq | Double negative B cells (DN2) |
| GSE118253 | GSM3323583 | CNTL | ATAC-seq | Resting Naive B cells (rN) |
| GSE118253 | GSM3323584 | CNTL | ATAC-seq | Switched Memory B cells (SM) |
| GSE118253 | GSM3323585 | CNTL | ATAC-seq | Transitional 3 B cells (T3) |
| GSE118253 | GSM3323586 | CNTL | ATAC-seq | Double negative B cells (DN2) |
| GSE118253 | GSM3323588 | CNTL | ATAC-seq | Resting Naive B cells (rN) |
| GSE118253 | GSM3323589 | CNTL | ATAC-seq | Activated Naive B cells (aN) |
| GSE118253 | GSM3323590 | CNTL | ATAC-seq | Double negative B cells (DN2) |
| GSE118253 | GSM3323592 | CNTL | ATAC-seq | Resting Naive B cells (rN) |
| GSE118253 | GSM3323593 | CNTL | ATAC-seq | Switched Memory B cells (SM) |
| GSE118253 | GSM3323594 | CNTL | ATAC-seq | Transitional 3 B cells (T3) |
| GSE118253 | GSM3323596 | CNTL | ATAC-seq | Resting Naive B cells (rN) |
| GSE118253 | GSM3323597 | CNTL | ATAC-seq | Transitional 3 B cells (T3) |
| GSE118253 | GSM3323598 | CNTL | ATAC-seq | Double negative B cells (DN2) |
| GSE118253 | GSM3323600 | CNTL | ATAC-seq | Switched Memory B cells (SM) |
| GSE118253 | GSM3323601 | CNTL | ATAC-seq | Transitional 3 B cells (T3) |
| GSE118253 | GSM3323602 | CNTL | ATAC-seq | Activated Naive B cells (aN) |
| GSE118253 | GSM3323604 | CNTL | ATAC-seq | Switched Memory B cells (SM) |
| GSE118253 | GSM3323605 | CNTL | ATAC-seq | Transitional 3 B cells (T3) |
| GSE118253 | GSM3323606 | CNTL | ATAC-seq | Double negative B cells (DN2) |
| GSE118253 | GSM3323608 | CNTL | ATAC-seq | Switched Memory B cells (SM) |
| GSE118253 | GSM3323609 | CNTL | ATAC-seq | Transitional 3 B cells (T3) |
| GSE118253 | GSM3323610 | CNTL | ATAC-seq | Activated Naive B cells (aN) |
| GSE118253 | GSM3323611 | CNTL | ATAC-seq | Double negative B cells (DN2) |
| GSE118253 | GSM3323613 | CNTL | ATAC-seq | Switched Memory B cells (SM) |
| GSE118253 | GSM3323614 | CNTL | ATAC-seq | Transitional 3 B cells (T3) |
| GSE118253 | GSM3323615 | SLE | ATAC-seq | Activated Naive B cells (aN) |
| GSE118253 | GSM3323616 | SLE | ATAC-seq | Double negative B cells (DN2) |
| GSE118253 | GSM3323617 | SLE | ATAC-seq | Switched Memory B cells (SM) |
| GSE118253 | GSM3323618 | SLE | ATAC-seq | Transitional 3 B cells (T3) |
| GSE118253 | GSM3323619 | SLE | ATAC-seq | Activated Naive B cells (aN) |
| GSE118253 | GSM3323620 | SLE | ATAC-seq | Double negative B cells (DN2) |
| GSE118253 | GSM3323621 | SLE | ATAC-seq | Switched Memory B cells (SM) |
| GSE118253 | GSM3323622 | SLE | ATAC-seq | Transitional 3 B cells (T3) |
| GSE118253 | GSM3323623 | SLE | ATAC-seq | Double negative B cells (DN2) |
| GSE118253 | GSM3323624 | SLE | ATAC-seq | Switched Memory B cells (SM) |
| GSE118253 | GSM3323625 | SLE | ATAC-seq | Activated Naive B cells (aN) |
| GSE118253 | GSM3323626 | SLE | ATAC-seq | Double negative B cells (DN2) |
| GSE118253 | GSM3323628 | SLE | ATAC-seq | Switched Memory B cells (SM) |
| GSE118253 | GSM3323629 | SLE | ATAC-seq | Transitional 3 B cells (T3) |
| GSE71338 | GSM2058522 | CNTL | ATAC-Seq | Naive B cell |
| GSE71338 | GSM2058523 | CNTL | ATAC-Seq | Naive B cell |
| GSE71338 | GSM2058524 | CNTL | ATAC-Seq | Naive B cell |
| GSE71338 | GSM2058525 | CNTL | ATAC-Seq | Naive B cell |
| GSE71338 | GSM2058526 | SLE | ATAC-Seq | Naive B cell |
| GSE71338 | GSM2058527 | SLE | ATAC-Seq | Naive B cell |
| GSE71338 | GSM2058528 | SLE | ATAC-Seq | Naive B cell |
| GSE71338 | GSM2058529 | SLE | ATAC-Seq | Naive B cell |
| GSE118254 | GSM3323630 | Control | RNA-Seq | Activated Naive B cells (aN) |
| GSE118254 | GSM3323631 | Control | RNA-Seq | Double negative B cells (DN2) |
| GSE118254 | GSM3323632 | Control | RNA-Seq | Switched Memory B cells (SM) |
| GSE118254 | GSM3323633 | Control | RNA-Seq | Transitional 3 B cells (T3) |
| GSE118254 | GSM3323634 | Control | RNA-Seq | Resting Naive B cells (rN) |
| GSE118254 | GSM3323635 | Control | RNA-Seq | Switched Memory B cells (SM) |
| GSE118254 | GSM3323636 | Control | RNA-Seq | Transitional 3 B cells (T3) |
| GSE118254 | GSM3323637 | Control | RNA-Seq | Double negative B cells (DN2) |
| GSE118254 | GSM3323638 | Control | RNA-Seq | Resting Naive B cells (rN) |
| GSE118254 | GSM3323639 | Control | RNA-Seq | Switched Memory B cells (SM) |
| GSE118254 | GSM3323640 | Control | RNA-Seq | Transitional 3 B cells (T3) |
| GSE118254 | GSM3323641 | Control | RNA-Seq | Activated Naive B cells (aN) |
| GSE118254 | GSM3323642 | Control | RNA-Seq | Double negative B cells (DN2) |
| GSE118254 | GSM3323644 | Control | RNA-Seq | Resting Naive B cells (rN) |
| GSE118254 | GSM3323645 | Control | RNA-Seq | Switched Memory B cells (SM) |
| GSE118254 | GSM3323646 | Control | RNA-Seq | Transitional 3 B cells (T3) |
| GSE118254 | GSM3323647 | Control | RNA-Seq | Activated Naive B cells (aN) |
| GSE118254 | GSM3323648 | Control | RNA-Seq | Switched Memory B cells (SM) |
| GSE118254 | GSM3323649 | Control | RNA-Seq | Transitional 3 B cells (T3) |
| GSE118254 | GSM3323650 | Control | RNA-Seq | Activated Naive B cells (aN) |
| GSE118254 | GSM3323651 | Control | RNA-Seq | Double negative B cells (DN2) |
| GSE118254 | GSM3323652 | Control | RNA-Seq | Resting Naive B cells (rN) |
| GSE118254 | GSM3323653 | Control | RNA-Seq | Switched Memory B cells (SM) |
| GSE118254 | GSM3323654 | Control | RNA-Seq | Transitional 3 B cells (T3) |
| GSE118254 | GSM3323655 | Control | RNA-Seq | Activated Naive B cells (aN) |
| GSE118254 | GSM3323656 | Control | RNA-Seq | Double negative B cells (DN2) |
| GSE118254 | GSM3323658 | Control | RNA-Seq | Resting Naive B cells (rN) |
| GSE118254 | GSM3323660 | Control | RNA-Seq | Transitional 3 B cells (T3) |
| GSE118254 | GSM3323659 | Control | RNA-Seq | Switched Memory B cells (SM) |
| GSE118254 | GSM3323661 | Control | RNA-Seq | Activated Naive B cells (aN) |
| GSE118254 | GSM3323662 | Control | RNA-Seq | Double negative B cells (DN2) |
| GSE118254 | GSM3323664 | Control | RNA-Seq | Resting Naive B cells (rN) |
| GSE118254 | GSM3323665 | Control | RNA-Seq | Switched Memory B cells (SM) |
| GSE118254 | GSM3323666 | Control | RNA-Seq | Transitional 3 B cells (T3) |
| GSE118254 | GSM3323667 | SLE | RNA-Seq | Activated Naive B cells (aN) |
| GSE118254 | GSM3323668 | SLE | RNA-Seq | Double negative B cells (DN2) |
| GSE118254 | GSM3323670 | SLE | RNA-Seq | Resting Naive B cells (rN) |
| GSE118254 | GSM3323671 | SLE | RNA-Seq | Switched Memory B cells (SM) |
| GSE118254 | GSM3323672 | SLE | RNA-Seq | Transitional 3 B cells (T3) |
| GSE118254 | GSM3323673 | SLE | RNA-Seq | Activated Naive B cells (aN) |
| GSE118254 | GSM3323674 | SLE | RNA-Seq | Double negative B cells (DN2) |
| GSE118254 | GSM3323675 | SLE | RNA-Seq | Resting Naive B cells (rN) |
| GSE118254 | GSM3323676 | SLE | RNA-Seq | Switched Memory B cells (SM) |
| GSE118254 | GSM3323677 | SLE | RNA-Seq | Transitional 3 B cells (T3) |
| GSE118254 | GSM3323678 | SLE | RNA-Seq | Activated Naive B cells (aN) |
| GSE118254 | GSM3323679 | SLE | RNA-Seq | Double negative B cells (DN2) |
| GSE118254 | GSM3323681 | SLE | RNA-Seq | Resting Naive B cells (rN) |
| GSE118254 | GSM3323682 | SLE | RNA-Seq | Switched Memory B cells (SM) |
| GSE118254 | GSM3323683 | SLE | RNA-Seq | Transitional 3 B cells (T3) |
| GSE118254 | GSM3323684 | SLE | RNA-Seq | Activated Naive B cells (aN) |
| GSE118254 | GSM3323685 | SLE | RNA-Seq | Double negative B cells (DN2) |
| GSE118254 | GSM3323688 | SLE | RNA-Seq | Switched Memory B cells (SM) |
| GSE118254 | GSM3323687 | SLE | RNA-Seq | Resting Naive B cells (rN) |
| GSE118254 | GSM3323689 | SLE | RNA-Seq | Transitional 3 B cells (T3) |
| GSE118254 | GSM3323690 | SLE | RNA-Seq | Activated Naive B cells (aN) |
| GSE118254 | GSM3323691 | SLE | RNA-Seq | Double negative B cells (DN2) |
| GSE118254 | GSM3323692 | SLE | RNA-Seq | Resting Naive B cells (rN) |
| GSE118254 | GSM3323694 | SLE | RNA-Seq | Transitional 3 B cells (T3) |
| GSE118254 | GSM3323693 | SLE | RNA-Seq | Switched Memory B cells (SM) |
| GSE118254 | GSM3323695 | SLE | RNA-Seq | Switched Memory B cells (SM) |
| GSE118254 | GSM3323696 | SLE | RNA-Seq | Transitional 3 B cells (T3) |
| GSE118254 | GSM3323697 | SLE | RNA-Seq | Activated Naive B cells (aN) |
| GSE118254 | GSM3323699 | SLE | RNA-Seq | Resting Naive B cells (rN) |
| GSE118254 | GSM3323698 | SLE | RNA-Seq | Double negative B cells (DN2) |
| GSE118254 | GSM3323700 | SLE | RNA-Seq | Switched Memory B cells (SM) |
| GSE118254 | GSM3323701 | SLE | RNA-Seq | Transitional 3 B cells (T3) |
| GSE118254 | GSM3323702 | SLE | RNA-Seq | Double negative B cells (DN2) |
| GSE118254 | GSM3323704 | SLE | RNA-Seq | Switched Memory B cells (SM) |
| GSE118254 | GSM3323703 | SLE | RNA-Seq | Resting Naive B cells (rN) |
| GSE118254 | GSM3323705 | SLE | RNA-Seq | Transitional 3 B cells (T3) |
| GSE118254 | GSM3323706 | SLE | RNA-Seq | Resting Naive B cells (rN) |
| GSE118254 | GSM3323707 | SLE | RNA-Seq | Activated Naive B cells (aN) |
| GSE118254 | GSM3323708 | SLE | RNA-Seq | Double negative B cells (DN2) |
| GSE118254 | GSM3323710 | SLE | RNA-Seq | Resting Naive B cells (rN) |
| GSE118254 | GSM3323711 | SLE | RNA-Seq | Switched Memory B cells (SM) |
| GSE118254 | GSM3323712 | SLE | RNA-Seq | Transitional 3 B cells (T3) |
| GSE92387 | GSM2428862 | SLE | RNA-Seq | Naïve B cell |
| GSE92387 | GSM2428863 | SLE | RNA-Seq | Switched memory |
| GSE92387 | GSM2428864 | SLE | RNA-Seq | Double negative 1 |
| GSE92387 | GSM2428865 | SLE | RNA-Seq | Double negative 2 |
| GSE92387 | GSM2428866 | SLE | RNA-Seq | Naïve |
| GSE92387 | GSM2428867 | SLE | RNA-Seq | Switched memory |
| GSE92387 | GSM2428868 | SLE | RNA-Seq | Double negative 1 |
| GSE92387 | GSM2428869 | SLE | RNA-Seq | Double negative 2 |
| GSE92387 | GSM2428870 | SLE | RNA-Seq | Naïve |
| GSE92387 | GSM2428871 | SLE | RNA-Seq | Switched memory |
| GSE92387 | GSM2428872 | SLE | RNA-Seq | Double negative 1 |
| GSE92387 | GSM2428873 | SLE | RNA-Seq | Double negative 2 |
| GSE92387 | GSM2428874 | CNTL | RNA-Seq | Naïve |
| GSE92387 | GSM2428875 | CNTL | RNA-Seq | Switched memory |
| GSE92387 | GSM2428876 | CNTL | RNA-Seq | Double negative 1 |
| GSE92387 | GSM2428877 | CNTL | RNA-Seq | Double negative 2 |
| GSE92387 | GSM2428878 | CNTL | RNA-Seq | Naïve |
| GSE92387 | GSM2428879 | CNTL | RNA-Seq | Switched memory |
| GSE92387 | GSM2428880 | CNTL | RNA-Seq | Double negative 1 |
| GSE92387 | GSM2428881 | CNTL | RNA-Seq | Double negative 2 |
| GSE92387 | GSM2428882 | CNTL | RNA-Seq | Naïve |
| GSE92387 | GSM2428883 | CNTL | RNA-Seq | Switched memory |
| GSE92387 | GSM2428884 | CNTL | RNA-Seq | Double negative 1 |
| GSE92387 | GSM2428885 | CNTL | RNA-Seq | Double negative 2 |
| GSE139359 | GSM4138735 | CNTL | ATAC-seq | NDN |
| GSE139359 | GSM4138736 | CNTL | ATAC-seq | NDN |
| GSE139359 | GSM4138737 | CNTL | ATAC-seq | NDN |
| GSE139359 | GSM4138738 | CNTL | ATAC-seq | NDN |
| GSE139359 | GSM4138739 | CNTL | ATAC-seq | NDN |
| GSE139359 | GSM4138740 | CNTL | ATAC-seq | NDN |
| GSE139359 | GSM4138741 | CNTL | ATAC-seq | NDN |
| GSE139359 | GSM4138742 | CNTL | ATAC-seq | NDN |
| GSE139359 | GSM4138743 | SLE | ATAC-seq | NDN |
| GSE139359 | GSM4138744 | SLE | ATAC-seq | NDN |
| GSE139359 | GSM4138745 | SLE | ATAC-seq | NDN |
| GSE139359 | GSM4138746 | SLE | ATAC-seq | NDN |
| GSE139359 | GSM4138747 | SLE | ATAC-seq | NDN |
| GSE139359 | GSM4138748 | SLE | ATAC-seq | NDN |
| GSE139359 | GSM4138749 | SLE | ATAC-seq | NDN |
| GSE139359 | GSM4138750 | SLE | ATAC-seq | NDN |
| GSE139359 | GSM4138751 | SLE | ATAC-seq | LDG |
| GSE139359 | GSM4138752 | SLE | ATAC-seq | LDG |
| GSE139359 | GSM4138753 | SLE | ATAC-seq | LDG |
| GSE139359 | GSM4138754 | SLE | ATAC-seq | LDG |
| GSE139359 | GSM4138755 | SLE | ATAC-seq | LDG |
| GSE139359 | GSM4138756 | SLE | ATAC-seq | LDG |
| GSE139359 | GSM4138757 | SLE | ATAC-seq | LDG |
| GSE139359 | GSM4138758 | SLE | ATAC-seq | LDG |
| GSE110017 | GSM2976426 | SLE | ATAC-seq | CD4+ T (PD1hi) |
| GSE110017 | GSM2976427 | SLE | ATAC-seq | CD4+ T (Tfh) |
| GSE110017 | GSM2976428 | SLE | ATAC-seq | CD4+ T (PD1hi) |
| GSE110017 | GSM2976429 | SLE | ATAC-seq | CD4+ T (PD1hi) |
| GSE110017 | GSM2976430 | SLE | ATAC-seq | CD4+ T (Tfh) |
| GSE110017 | GSM2976431 | SLE | ATAC-seq | CD4+ T (PD1hi) |
| GSE118255 | GSM3323713 | CNTL | RRBS | Activated Naive B cells (aN) |
| GSE118255 | GSM3323714 | CNTL | RRBS | Double negative B cells (DN2) |
| GSE118255 | GSM3323715 | CNTL | RRBS | Resting Naive B cells (rN) |
| GSE118255 | GSM3323716 | CNTL | RRBS | Switched Memory B cells (SM) |
| GSE118255 | GSM3323717 | CNTL | RRBS | Transitional 3 B cells (T3) |
| GSE118255 | GSM3323718 | CNTL | RRBS | Activated Naive B cells (aN) |
| GSE118255 | GSM3323719 | CNTL | RRBS | Double negative B cells (DN2) |
| GSE118255 | GSM3323720 | CNTL | RRBS | Resting Naive B cells (rN) |
| GSE118255 | GSM3323721 | CNTL | RRBS | Switched Memory B cells (SM) |
| GSE118255 | GSM3323722 | CNTL | RRBS | Transitional 3 B cells (T3) |
| GSE118255 | GSM3323723 | CNTL | RRBS | Activated Naive B cells (aN) |
| GSE118255 | GSM3323724 | CNTL | RRBS | Double negative B cells (DN2) |
| GSE118255 | GSM3323725 | CNTL | RRBS | Resting Naive B cells (rN) |
| GSE118255 | GSM3323726 | CNTL | RRBS | Switched Memory B cells (SM) |
| GSE118255 | GSM3323727 | CNTL | RRBS | Transitional 3 B cells (T3) |
| GSE118255 | GSM3323728 | CNTL | RRBS | Activated Naive B cells (aN) |
| GSE118255 | GSM3323729 | CNTL | RRBS | Double negative B cells (DN2) |
| GSE118255 | GSM3323730 | CNTL | RRBS | Resting Naive B cells (rN) |
| GSE118255 | GSM3323731 | CNTL | RRBS | Switched Memory B cells (SM) |
| GSE118255 | GSM3323732 | CNTL | RRBS | Transitional 3 B cells (T3) |
| GSE118255 | GSM3323733 | CNTL | RRBS | Activated Naive B cells (aN) |
| GSE118255 | GSM3323734 | CNTL | RRBS | Double negative B cells (DN2) |
| GSE118255 | GSM3323736 | CNTL | RRBS | Resting Naive B cells (rN) |
| GSE118255 | GSM3323737 | CNTL | RRBS | Switched Memory B cells (SM) |
| GSE118255 | GSM3323738 | CNTL | RRBS | Transitional 3 B cells (T3) |
| GSE118255 | GSM3323739 | CNTL | RRBS | Activated Naive B cells (aN) |
| GSE118255 | GSM3323740 | CNTL | RRBS | Double negative B cells (DN2) |
| GSE118255 | GSM3323741 | CNTL | RRBS | Resting Naive B cells (rN) |
| GSE118255 | GSM3323742 | CNTL | RRBS | Switched Memory B cells (SM) |
| GSE118255 | GSM3323743 | CNTL | RRBS | Transitional 3 B cells (T3) |
| GSE118255 | GSM3323744 | CNTL | RRBS | Activated Naive B cells (aN) |
| GSE118255 | GSM3323745 | CNTL | RRBS | Double negative B cells (DN2) |
| GSE118255 | GSM3323747 | CNTL | RRBS | Resting Naive B cells (rN) |
| GSE118255 | GSM3323748 | CNTL | RRBS | Switched Memory B cells (SM) |
| GSE118255 | GSM3323749 | CNTL | RRBS | Transitional 3 B cells (T3) |
| GSE118255 | GSM3323750 | CNTL | RRBS | Activated Naive B cells (aN) |
| GSE118255 | GSM3323751 | CNTL | RRBS | Double negative B cells (DN2) |
| GSE118255 | GSM3323753 | CNTL | RRBS | Resting Naive B cells (rN) |
| GSE118255 | GSM3323754 | CNTL | RRBS | Switched Memory B cells (SM) |
| GSE118255 | GSM3323755 | CNTL | RRBS | Transitional 3 B cells (T3) |
| GSE118255 | GSM3323756 | SLE | RRBS | Activated Naive B cells (aN) |
| GSE118255 | GSM3323757 | SLE | RRBS | Double negative B cells (DN2) |
| GSE118255 | GSM3323758 | SLE | RRBS | Resting Naive B cells (rN) |
| GSE118255 | GSM3323759 | SLE | RRBS | Switched Memory B cells (SM) |
| GSE118255 | GSM3323760 | SLE | RRBS | Transitional 3 B cells (T3) |
| GSE118255 | GSM3323761 | SLE | RRBS | Activated Naive B cells (aN) |
| GSE118255 | GSM3323762 | SLE | RRBS | Double negative B cells (DN2) |
| GSE118255 | GSM3323763 | SLE | RRBS | Resting Naive B cells (rN) |
| GSE118255 | GSM3323764 | SLE | RRBS | Switched Memory B cells (SM) |
| GSE118255 | GSM3323765 | SLE | RRBS | Transitional 3 B cells (T3) |
| GSE118255 | GSM3323766 | SLE | RRBS | Activated Naive B cells (aN) |
| GSE118255 | GSM3323767 | SLE | RRBS | Double negative B cells (DN2) |
| GSE118255 | GSM3323768 | SLE | RRBS | Resting Naive B cells (rN) |
| GSE118255 | GSM3323769 | SLE | RRBS | Switched Memory B cells (SM) |
| GSE118255 | GSM3323770 | SLE | RRBS | Transitional 3 B cells (T3) |
| GSE118255 | GSM3323771 | SLE | RRBS | Activated Naive B cells (aN) |
| GSE118255 | GSM3323772 | SLE | RRBS | Double negative B cells (DN2) |
| GSE118255 | GSM3323773 | SLE | RRBS | Resting Naive B cells (rN) |
| GSE118255 | GSM3323774 | SLE | RRBS | Switched Memory B cells (SM) |
| GSE118255 | GSM3323775 | SLE | RRBS | Transitional 3 B cells (T3) |
| GSE118255 | GSM3323776 | SLE | RRBS | Activated Naive B cells (aN) |
| GSE118255 | GSM3323777 | SLE | RRBS | Double negative B cells (DN2) |
| GSE118255 | GSM3323779 | SLE | RRBS | Resting Naive B cells (rN) |
| GSE118255 | GSM3323780 | SLE | RRBS | Switched Memory B cells (SM) |
| GSE118255 | GSM3323781 | SLE | RRBS | Transitional 3 B cells (T3) |
| GSE118255 | GSM3323782 | SLE | RRBS | Activated Naive B cells (aN) |
| GSE118255 | GSM3323783 | SLE | RRBS | Double negative B cells (DN2) |
| GSE118255 | GSM3323784 | SLE | RRBS | Resting Naive B cells (rN) |
| GSE118255 | GSM3323785 | SLE | RRBS | Switched Memory B cells (SM) |
| GSE118255 | GSM3323786 | SLE | RRBS | Transitional 3 B cells (T3) |
| GSE118255 | GSM3323787 | SLE | RRBS | Activated Naive B cells (aN) |
| GSE118255 | GSM3323788 | SLE | RRBS | Double negative B cells (DN2) |
| GSE118255 | GSM3323790 | SLE | RRBS | Resting Naive B cells (rN) |
| GSE118255 | GSM3323791 | SLE | RRBS | Switched Memory B cells (SM) |
| GSE118255 | GSM3323792 | SLE | RRBS | Transitional 3 B cells (T3) |
| GSE118255 | GSM3323793 | SLE | RRBS | Activated Naive B cells (aN) |
| GSE118255 | GSM3323794 | SLE | RRBS | Double negative B cells (DN2) |
| GSE118255 | GSM3323796 | SLE | RRBS | Resting Naive B cells (rN) |
| GSE118255 | GSM3323797 | SLE | RRBS | Switched Memory B cells (SM) |
| GSE118255 | GSM3323798 | SLE | RRBS | Transitional 3 B cells (T3) |
| GSE118255 | GSM3323799 | SLE | RRBS | Activated Naive B cells (aN) |
| GSE118255 | GSM3323800 | SLE | RRBS | Double negative B cells (DN2) |
| GSE118255 | GSM3323802 | SLE | RRBS | Resting Naive B cells (rN) |
| GSE118255 | GSM3323803 | SLE | RRBS | Switched Memory B cells (SM) |
| GSE118255 | GSM3323804 | SLE | RRBS | Transitional 3 B cells (T3) |

**Table S3:**
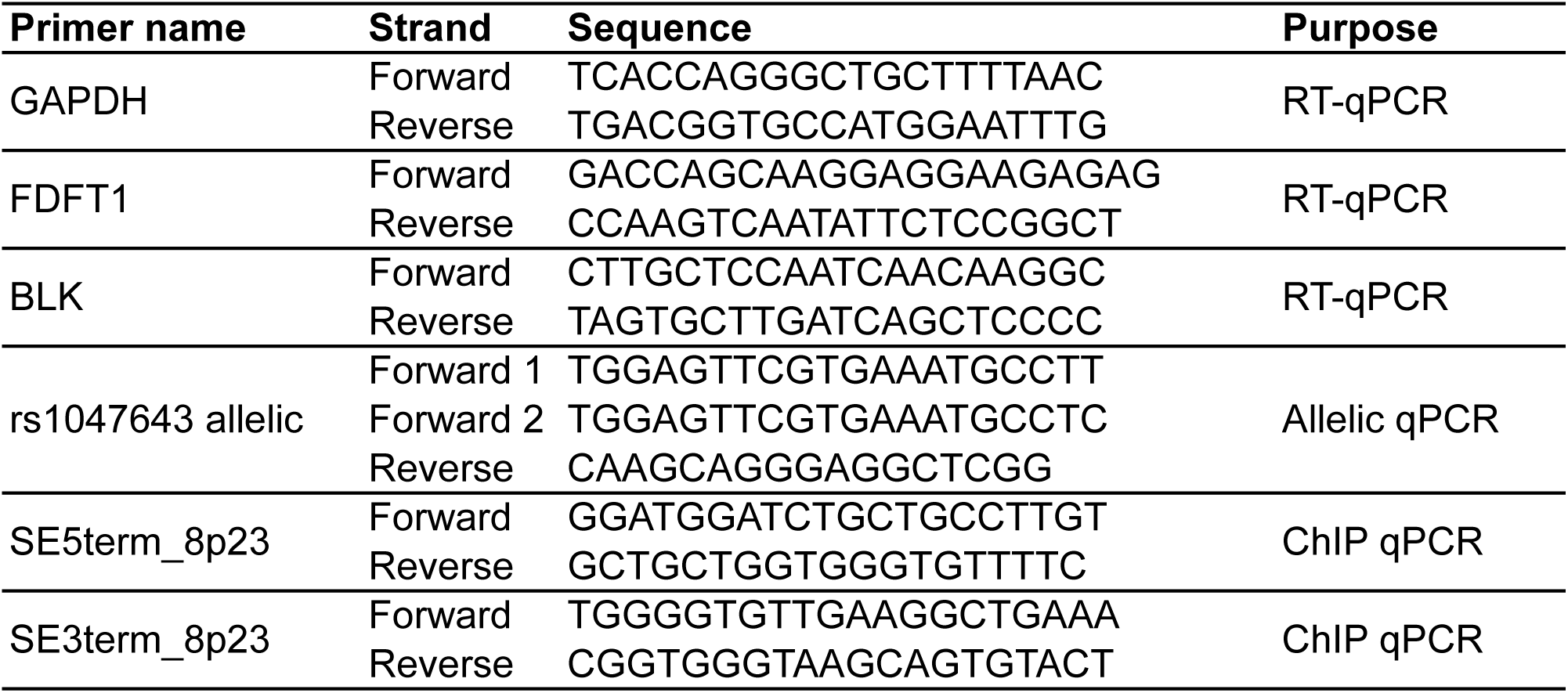
List of primers used in this study.

## Notes

### Competing Interest Statement

The authors have declared no competing interest.

## References

1. Parker SCJ, Stitzel ML, Taylor DL, Orozco JM, Erdos MR, Akiyama JA, et al. Chromatin stretch enhancer states drive cell-specific gene regulation and harbor human disease risk variants. Proceedings of the National Academy of Sciences. 2013;110(44):17921–6.

2. Whyte WA, Orlando DA, Hnisz D, Abraham BJ, Lin CY, Kagey MH, et al. Master Transcription Factors and Mediator Establish Super-Enhancers at Key Cell Identity Genes. Cell. 2013;153(2):307–19.

3. Vahedi G, Kanno Y, Furumoto Y, Jiang K, Parker SCJ, Erdos MR, et al. Super-enhancers delineate disease-associated regulatory nodes in T cells. Nature. 2015;520(7548):558–62.

4. Hnisz D, Abraham Brian J, Lee Tong I, Lau A, Saint-André V, Sigova Alla A, et al. Super-Enhancers in the Control of Cell Identity and Disease. Cell. 2013;155(4):934–47.

5. Decker T, Kovarik P. Serine phosphorylation of STATs. Oncogene. 2000;19(21):2628–37.

6. Levy DE, Darnell JE. STATs: transcriptional control and biological impact. Nature Reviews Molecular Cell Biology. 2002;3(9):651–62.

7. Yu H, Pardoll D, Jove R. STATs in cancer inflammation and immunity: a leading role for STAT3. Nature Reviews Cancer. 2009;9(11):798–809.

8. Rahman A, Isenberg DA. Systemic Lupus Erythematosus. New England Journal of Medicine. 2008;358(9):929–39.

9. Catalina MD, Owen KA, Labonte AC, Grammer AC, Lipsky PE. The pathogenesis of systemic lupus erythematosus: Harnessing big data to understand the molecular basis of lupus. Journal of Autoimmunity. 2020;110:102359.

10. Yin X, Kim K, Suetsugu H, Bang S-Y, Wen L, Koido M, et al. Meta-analysis of 208370 East Asians identifies 113 susceptibility loci for systemic lupus erythematosus. Annals of the Rheumatic Diseases. 2021;80(5):632–40.

11. Guthridge JM, Lu R, Sun H, Sun C, Wiley GB, Dominguez N, et al. Two functional lupus-associated BLK promoter variants control cell-type- and developmental-stage-specific transcription. American journal of human genetics. 2014;94(4):586–98.

12. Sun C, Molineros JE, Looger LL, Zhou X-j, Kim K, Okada Y, et al. High-density genotyping of immune-related loci identifies new SLE risk variants in individuals with Asian ancestry. Nature Genetics. 2016;48(3):323–30.

13. Morris DL, Sheng Y, Zhang Y, Wang Y-F, Zhu Z, Tombleson P, et al. Genome-wide association meta-analysis in Chinese and European individuals identifies ten new loci associated with systemic lupus erythematosus. Nature Genetics. 2016;48(8):940–6.

14. Gallagher MD, Chen-Plotkin AS. The Post-GWAS Era: From Association to Function. American journal of human genetics. 2018;102(5):717–30.

15. Pastinen T, Hudson TJ. Cis-Acting Regulatory Variation in the Human Genome. Science. 2004;306(5696):647–50.

16. Yan H, Yuan W, Velculescu VE, Vogelstein B, Kinzler KW. Allelic variation in human gene expression. Science. 2002;297(5584):1143.

17. Li Q, Seo JH, Stranger B, McKenna A, Pe’er I, Laframboise T, et al. Integrative eQTL-based analyses reveal the biology of breast cancer risk loci. Cell. 2013;152(3):633–41.

18. Zhang S, Zhang H, Zhou Y, Qiao M, Zhao S, Kozlova A, et al. Allele-specific open chromatin in human iPSC neurons elucidates functional disease variants. Science. 2020;369(6503):561–5.

19. Pollard KS, Serre D, Wang X, Tao H, Grundberg E, Hudson TJ, et al. A genome-wide approach to identifying novel-imprinted genes. Human genetics. 2008;122(6):625–34.

20. Zhang Y, Li X, Gibson A, Edberg J, Kimberly RP, Absher DM. Skewed allelic expression on X chromosome associated with aberrant expression of XIST on systemic lupus erythematosus lymphocytes. Human molecular genetics. 2020;29(15):2523–34.

21. Zhang Y, Wagner EK, Guo X, May I, Cai Q, Zheng W, et al. Long intergenic non-coding RNA expression signature in human breast cancer. Scientific reports. 2016;6:37821.

22. Kim D, Paggi JM, Park C, Bennett C, Salzberg SL. Graph-based genome alignment and genotyping with HISAT2 and HISAT-genotype. Nature biotechnology. 2019;37(8):907–15.

23. Li H, Handsaker B, Wysoker A, Fennell T, Ruan J, Homer N, et al. The Sequence Alignment/Map format and SAMtools. Bioinformatics. 2009;25(16):2078–9.

24. Zhang Y, Delahanty R, Guo X, Zheng W, Long J. Integrative genomic analysis reveals functional diversification of APOBEC gene family in breast cancer. Human Genomics. 2015;9(1):34.

25. Langmead B, Salzberg SL. Fast gapped-read alignment with Bowtie 2. Nature methods. 2012;9(4):357–9.

26. Das S, Forer L, Schonherr S, Sidore C, Locke AE, Kwong A, et al. Next-generation genotype imputation service and methods. Nat Genet. 2016;48(10):1284–7.

27. Shi J, Zhang Y, Zheng W, Michailidou K, Ghoussaini M, Bolla MK, et al. Fine-scale mapping of 8q24 locus identifies multiple independent risk variants for breast cancer. Int J Cancer. 2016;139(6):1303–17.

28. Parker SC, Stitzel ML, Taylor DL, Orozco JM, Erdos MR, Akiyama JA, et al. Chromatin stretch enhancer states drive cell-specific gene regulation and harbor human disease risk variants. Proceedings of the National Academy of Sciences of the United States of America. 2013;110(44):17921–6.

29. Consortium GT, Laboratory DA, Coordinating Center -Analysis Working G, Statistical Methods groups-Analysis Working G, Enhancing Gg, Fund NIHC, et al. Genetic effects on gene expression across human tissues. Nature. 2017;550(7675):204–13.

30. Ward LD, Kellis M. HaploReg v4: systematic mining of putative causal variants, cell types, regulators and target genes for human complex traits and disease. Nucleic acids research. 2016;44(D1):D877–81.

31. Westra H-J, Peters MJ, Esko T, Yaghootkar H, Schurmann C, Kettunen J, et al. Systematic identification of trans eQTLs as putative drivers of known disease associations. Nature Genetics. 2013;45(10):1238–43.

32. Rao Suhas SP, Huntley Miriam H, Durand Neva C, Stamenova Elena K, Bochkov Ivan D, Robinson James T, et al. A 3D Map of the Human Genome at Kilobase Resolution Reveals Principles of Chromatin Looping. Cell. 2014;159(7):1665–80.

33. Wingett S, Ewels P, Furlan-Magaril M, Nagano T, Schoenfelder S, Fraser P, et al. HiCUP: pipeline for mapping and processing Hi-C data. F1000Research. 2015;4(1310).

34. Heinz S, Benner C, Spann N, Bertolino E, Lin YC, Laslo P, et al. Simple Combinations of Lineage-Determining Transcription Factors Prime *cis*-Regulatory Elements Required for Macrophage and B Cell Identities. Molecular Cell. 2010;38(4):576–89.

35. Scharer CD, Blalock EL, Mi T, Barwick BG, Jenks SA, Deguchi T, et al. Epigenetic programming underpins B cell dysfunction in human SLE. Nature Immunology. 2019;20(8):1071–82.

36. Madoux F, Koenig M, Nelson E, Chowdhury S, Cameron M, Mercer B, et al. Modulators of STAT Transcription Factors for the Targeted Therapy of Cancer (STAT3 Activators). Bethesda: National Center for Biotechnology Information; 2010.

37. Saijo K, Schmedt C, Su Ih, Karasuyama H, Lowell CA, Reth M, et al. Essential role of Src-family protein tyrosine kinases in NF-κB activation during B cell development. Nature Immunology. 2003;4(3):274–9.

38. Van Ness B, Ramos C, Haznadar M, Hoering A, Haessler J, Crowley J, et al. Genomic variation in myeloma: design, content, and initial application of the Bank On A Cure SNP Panel to detect associations with progression-free survival. BMC medicine. 2008;6:26.

39. Skibola CF, Bracci PM, Halperin E, Nieters A, Hubbard A, Paynter RA, et al. Polymorphisms in the Estrogen Receptor 1 and Vitamin C and Matrix Metalloproteinase Gene Families Are Associated with Susceptibility to Lymphoma. PLOS ONE. 2008;3(7):e2816.

40. Huang X, Meng B, Iqbal J, Ding BB, Perry AM, Cao W, et al. Activation of the STAT3 signaling pathway is associated with poor survival in diffuse large B-cell lymphoma treated with R-CHOP. Journal of clinical oncology. 2013;31(36):4520–8.

41. Jung S-H, Ahn S-Y, Choi H-W, Shin M-G, Lee S-S, Yang D-H, et al. STAT3 expression is associated with poor survival in non-elderly adult patients with newly diagnosed multiple myeloma. Blood Res. 2017;52(4):293–9.

42. Avery DT, Deenick EK, Ma CS, Suryani S, Simpson N, Chew GY, et al. B cell– intrinsic signaling through IL-21 receptor and STAT3 is required for establishing long-lived antibody responses in humans. Journal of Experimental Medicine. 2010;207(1):155–71.

43. Ding C, Chen X, Dascani P, Hu X, Bolli R, Zhang H-g, et al. STAT3 Signaling in B Cells Is Critical for Germinal Center Maintenance and Contributes to the Pathogenesis of Murine Models of Lupus. The Journal of Immunology. 2016;196(11):4477–86.

44. Demirci FY, Wang X, Morris DL, Feingold E, Bernatsky S, Pineau C, et al. Multiple signals at the extended 8p23 locus are associated with susceptibility to systemic lupus erythematosus. Journal of Medical Genetics. 2017;54(6):381–9.

45. Tozawa R-i, Ishibashi S, Osuga J-i, Yagyu H, Oka T, Chen Z, et al. Embryonic Lethality and Defective Neural Tube Closure in Mice Lacking Squalene Synthase. Journal of Biological Chemistry. 1999;274(43):30843–8.

46. Tisseverasinghe A, Lim S, Greenwood C, Urowitz M, Gladman D, Fortin PR. Association between serum total cholesterol level and renal outcome in systemic lupus erythematosus. Arthritis & Rheumatism. 2006;54(7):2211–9.

